# Hilar mossy cells control structural and functional organization in the dentate gyrus

**DOI:** 10.64898/2026.07.15.738260

**Authors:** Corwin R. Butler, Connor R. Squellati, Alyssa B. Danis, Arielle Isakharov, Gary L. Westbrook, Eric Schnell

**Author notes:** **Corresponding author:** Eric Schnell MD, PhD, Portland VA Healthcare System, 3710 SW US Veteran’s Hospital Rd, P3ANES, Portland, OR, USA, 97239. The authors declare no competing financial interests.

## Abstract

Hilar mossy cells in the dentate gyrus project widely throughout the hippocampus, broadly contributing to circuit function. Their loss in disease is associated with local functional and structural rearrangements, including retrograde granule cell axon sprouting, aberrant neurogenesis, and disinhibition. To examine how mossy cell loss contributes to these circuit rearrangements, we ablated or silenced hilar mossy cells using viral approaches in transgenic (Crlr-Cre) mice. Both mossy cell ablation and silencing dramatically altered dentate gyrus structure and function, as assessed using immunohistochemical, viral labeling, electrophysiology, and anatomical methods. Both manipulations accelerated the maturation of adult-born neurons, but did not alter neuroblast proliferation or cause granule cell axon sprouting. However, mossy cell ablation, but not silencing, caused collapse of the inner molecular layer accompanied by proximal translocation of middle molecular layer inputs. In both cases, granule cell activity measured by cFos labeling and seizure susceptibility were unchanged after mossy cell loss, indicating functional compensation for the altered network organization. Our results highlight how mossy cells influence dentate gyrus organization and adult neurogenesis but also demonstrate the resilience of the hippocampal circuit to structural or functional perturbations.

## Introduction

The dentate gyrus is classically considered a “gate” controlling hippocampal circuit activity. The principal neurons, dentate granule cells (DGCs), have apical dendrites that traverse three distinct laminar excitatory axon tracts within the molecular layer, and fire sparsely as a result of cell intrinsic and network characteristics (Hjorth-Simonsen, 1972; Treves and Rolls, 1994; Buckmaster et al., 1996; Witter, 2007; Barth and Poulet, 2012). Mossy cells, excitatory interneurons in the dentate hilus, innervate granule cells in the inner molecular layer (IML). Mossy cells project bilaterally and widely throughout the dorsoventral axis of the hippocampus and drive strong feedforward inhibition, which has led to hypotheses that these neurons broadly coordinate hippocampal function including pattern separation and regulation of network excitability (Scharfman, 2016; Galloni et al., 2022).

Mossy cell loss after traumatic brain injury and in epilepsy (Babb et al., 1984; De Lanerolle et al., 1989; Buckmaster and Dudek, 1997; Blümcke et al., 2000; Santhakumar et al., 2000) correlates with functional and structural changes in the hippocampus, such as aberrant neurogenesis and retrograde granule cell axon (mossy fiber) sprouting (De Lanerolle et al., 1989; Lowenstein et al., 1992; Blümcke et al., 2000; Overstreet-Wadiche et al., 2006; Bui et al., 2018; Koyama and Ikegaya, 2018). Thus, it has been suggested that mossy cell loss drives these structural changes and contributes to circuit dysfunction (Scharfman, 1994; Sloviter et al., 2003; Jinde et al., 2012; Jinde et al., 2013; Botterill et al., 2019). For example, sprouted granule cell axon terminals occupy dendritic regions “vacated” following the loss of mossy cell axons, and the early glutamatergic innervation of immature adult-born DGCs by mossy cells positions them well to impact hippocampal adult neurogenesis (Overstreet-Wadiche et al., 2006; Chancey et al., 2014). However, a direct and causal relationship between mossy cell loss and associated circuit rearrangements in disease is difficult to establish because of concurrent metabolic and inflammatory changes (De Lanerolle et al., 1989; Ekdahl et al., 2003; Iosif et al., 2006; Ziv et al., 2006).

To address this issue, we combined a mossy cell-selective Cre-expressing mouse line, Crlr-Cre (Jinde et al., 2012), with Cre-dependent viral vectors to selectively eliminate or silence mossy cells in the adult mouse hippocampus. The selective removal of mossy cell inputs drove functional and structural changes in the dentate gyrus, including acceleration of adult-born DGC dendritic outgrowth and IML collapse. Despite these dramatic circuit alterations, however, granule cell activity and seizure susceptibility were grossly unchanged, demonstrating the remarkable ability of the hippocampal circuit to adapt following mossy cell loss in the absence of other associated disease processes.

## Results

### Virus-mediated ablation or permanent silencing of hilar mossy cells

The development of mouse lines to genetically target hilar mossy cells has greatly facilitated understanding of these neurons (Jinde et al., 2012; Gulfo et al., 2023). To assess the contributions of mossy cells to hippocampal circuit structure and function, we injected Cre-dependent caspase-expressing viral vectors into the hippocampi of mossy cell-selective Crlr-Cre mice to ablate hilar mossy cells.

To validate this approach, we examined tissue 6 weeks after virus injection, using anti-calretinin immunohistochemistry to identify hilar mossy cells and their axons (Fujise et al., 1998; Alcantara-Gonzalez et al., 2025). Following AAV5-flex-taCasp3 virus injection, ∼89% of calretinin-positive hilar cell bodies and ∼84% of glutamate ionotropic receptor AMPA type subunit 2 (GluA2)-positive hilar cell bodies were lost in Cre-positive mice, compared to Cre-negative (WT) mice that received identical virus injections, consistent with widespread removal of mossy cells across the septotemporal axis of the hippocampus (Figure 1A-C; Supplemental Figure 1). This mossy cell loss is comparable to inducible expression of the diphtheria toxin receptor in the same mouse line (Jinde et al., 2012). Although not a complete ablation, this value is substantially higher than chemoconvulsant models of epilepsy, which typically result in ∼40-75% mossy cell loss (Buckmaster and Dudek, 1997; Blümcke et al., 2000; Bui et al., 2018; Butler et al., 2022), and importantly, without the other chemoconvulsant-induced levels of circuit activation across the brain. Expression of Cre recombinase can occur in proximal CA3 pyramidal neurons in this mouse line (Jinde et al., 2012), but we observed no significant difference in CA3+ cell density following hilar AAV5-flex-taCasp3 injection (Supplemental Figure 2 A,B). Mossy cell ablation transiently increased microglial activation in the dentate hilus and inner molecular layer (IML) (Supplemental Figure 3A-C). To avoid the potential impact of microglial activation (Ekdahl et al., 2003; Iosif et al., 2006; Ekdahl et al., 2009), we delayed additional manipulations or assessments until 3 weeks after caspase virus injection, at which point glial marker expression had returned to basal levels (Supplemental Figure 3A-C).

**Figure 1:**
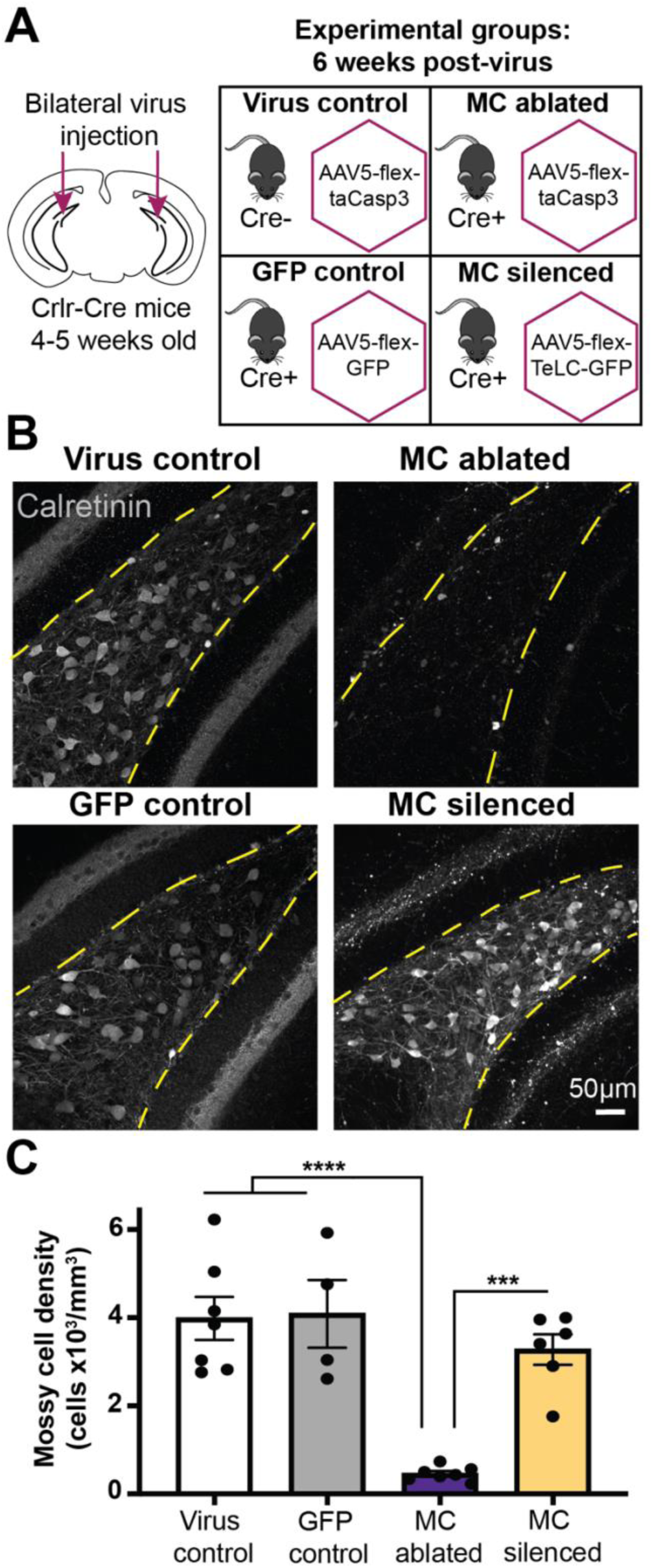
Hilar mossy cell targeting by viral vector injections *in vivo*. A) Schematic of AAV injection coordinates in Calcitonin receptor-like receptor (Crlr-Cre) mice, with description of experimental groups including mouse genotype and viral vector used. B) Representative images of dentate sections stained for the mossy cell (MC) marker calretinin in each condition, 6 weeks after virus injection, demonstrating hilar mossy cell loss in MC ablated mice and preserved mossy cells in MC silenced mice. Yellow dashed lines note hilus in images C) Summary data of hilar mossy cell densities for each group, 6 weeks after virus injection (Virus control n = 7 mice, GFP control n = 4 mice, MC ablated n = 7 mice, and MC silenced n = 6 mice; *** p <0.001, **** p <0.0001).

As an alternative to mossy cell ablation, we used Cre-dependent viral vectors to permanently silence mossy cell outputs with tetanus toxin (Murray et al., 2011). AAV-mediated expression of tetanus toxin light chain completely prevents synaptic transmission while preserving cell integrity and axonal projections, thus facilitating discrimination between loss of cells and loss of functional activity (Woods et al., 2018). In Crlr-Cre mice injected with Cre-dependent tetanus toxin light chain virus (AAV5-flex-TeLC-GFP), the density of hilar mossy cells was not altered relative to either WT mice or control virus (AAV5-flex-GFP)-injected mice, but was greater than mossy cell-ablated mice (Figure 1A-C). Viral targeting of calretinin-positive mossy cells was similarly high in both silenced and control mice (GFP= 89.9 ±2.1% of calretinin positive mossy cells, N=10 mice; TeLC-GFP= 90.7 ±1.1% of calretinin positive mossy cells, N=9 mice; Student’s t-test, F(9,8)=4.112, p=0.74). GFP-positive hilar neurons and axons were present throughout the dorsoventral hippocampus with either vector, including more dorsal sections where mossy cells have substantially less calretinin expression (Houser et al., 2020; Botterill et al., 2021). A subset of mice were examined to asses potential off-target virus expression, we observed very few GFP+ neurons in the proximal CA3 pyramidal cell layer (GFP control= 1.5 ±1.3%, N=3 mice; MC silenced= 2.9 ±1.4%, N= 3 mice), indicating very limited impact of these viral manipulations on CA3 circuits (Supplemental Figure 2 C,D). Unlike tissue after caspase-mediated mossy cell loss, tetanus toxin and GFP-expressing tissue did not exhibit microglial activation (Supplemental Figure 3D-F).

### Accelerated adult-born DGC dendritic maturation following mossy cell loss

The maturation of adult-born granule cells in mice follows a stereotyped rate and pattern of dendritic outgrowth and synapse acquisition (van Praag et al., 2002; Tashiro et al., 2006; Toni et al., 2008). Functional glutamatergic synaptogenesis critically guides this process (Tashiro et al., 2006; Schnell et al., 2014; Doengi et al., 2016), with mossy cells providing the first glutamatergic inputs to adult-born DGCs (Chancey et al., 2014). The rate and pattern of dendritic outgrowth by adult-born neurons is dramatically altered after brain injury (Overstreet-Wadiche et al., 2006; Parent et al., 2006; Niv et al., 2012; Villasana et al., 2015) and correlates with mossy cell loss (Overstreet-Wadiche et al., 2006). However, the association between mossy cell loss and aberrant neurogenesis could reflect changes secondary to other injury-related signaling.

We used GFP-expressing retroviral vectors (van Praag et al., 2002) to selectively label adult-born DGCs generated 3 weeks after mossy cell ablation. Mice were sacrificed 14 days after retroviral injection to identify immature adult-born DGCs at an intermediate stage of dendritic development, during which their glutamatergic inputs are exclusively mediated by mossy cell innervation (Chancey et al., 2014). Dendritic outgrowth was dramatically increased in 14-day old DGCs in the absence of mossy cell inputs, compared to control (virus-injected, Cre-negative) (Figure 2A-E). This was evident in total dendritic length, the number of dendritic branches per neuron, and in the number of branch points at distances closer to the soma (Figure 2C-E). We examined if this effect was associated with alterations in cellular proliferation or survival by labeling cells with the mitotic marker BrdU (at 24hrs or 21 days after i.p. administration) or the immature DGC marker doublecortin (DCX) (Brown et al., 2003) in a separate set of mice. We found no difference in the density of BrdU+ cells in either the subgranular zone or granule cell layer at 24hrs or 21 days post-administration for mice following mossy cell ablation (Supplemental Figure 4). There was also no difference in DCX+ cell density or outward migration following mossy cell ablation (Supplemental Figure 5A-C). Thus, the absence of mossy cell inputs directly altered dendritic outgrowth of adult-born hippocampal granule cells with no gross changes to the overall process of adult neurogenesis.

**Figure 2:**
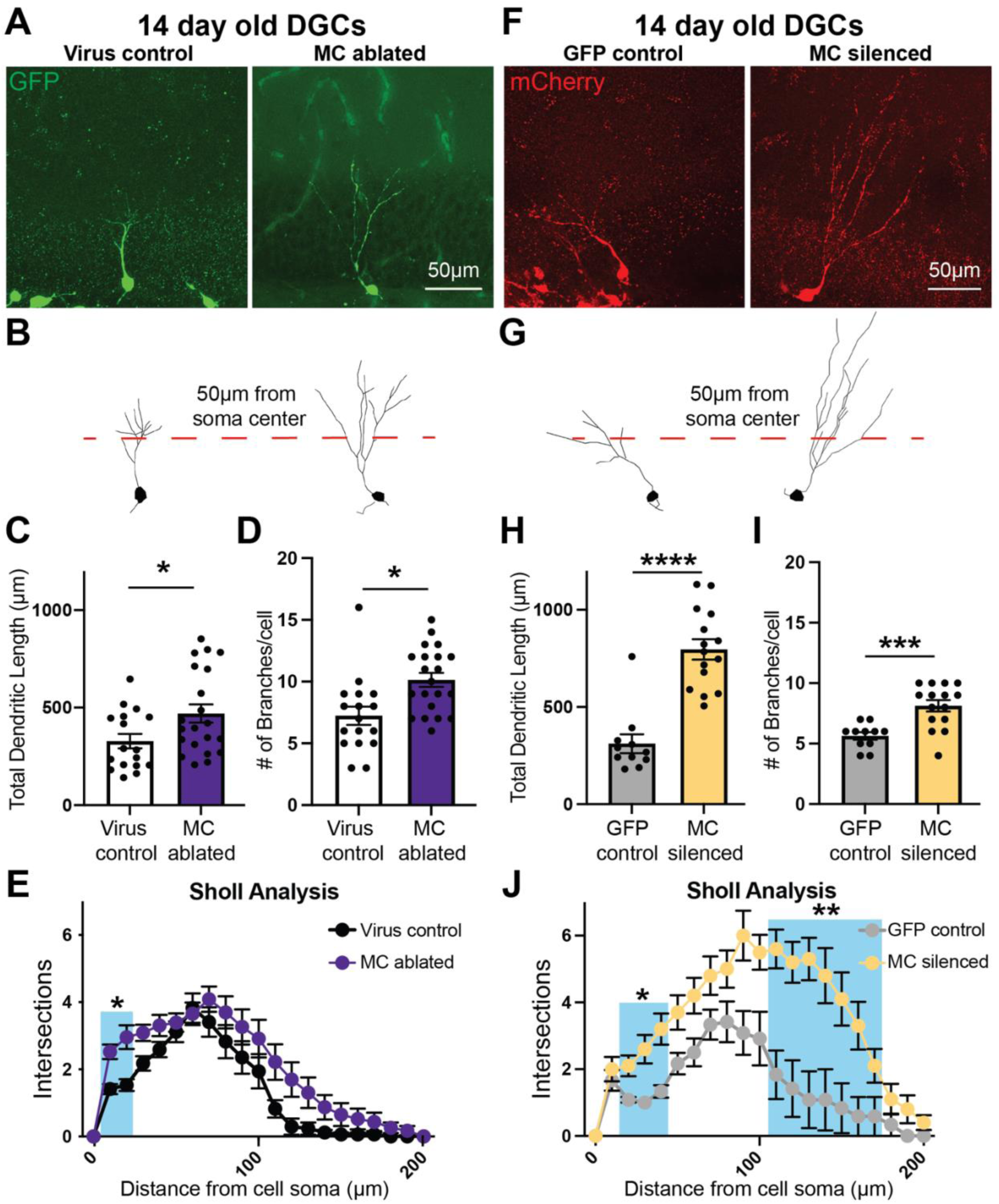
Enhanced dendritic growth of immature adult-born DGCs in the absence of mossy cell inputs. A) Representative GFP-expressing adult-born dentate granule cells (DGCs) in virus-injected control and mossy cell (MC) ablated mice, 14 days after retroviral labeling/mitosis. B) Skeleton tracing of cell morphology from the representative images in A, with a dashed line 50µm from cell body center. C&D) Total dendritic length and number of branches per cell of neurons from virus control and MC ablated mice (Virus control n = 17 cells/ 5 mice, MC ablated n = 21 cells/ 6 mice; * p <0.05). E) Sholl analysis demonstrates more extensive proximal dendritic arborization for DGCs born after mossy cell ablation (* p= 0.02 and 0.01 at intervals 10 and 20µm distance from soma, blue shading, all other points not significant). F) Representative mCherry-expressing adult-born (DGCs) in control (AAV5-flex-GFP) virus injected and MC silenced (AAV5-flex-TeLC-injected) mice, 14 days after retroviral labeling. G) Skeleton tracing of cell morphology for the representative images in G, with a dashed line 50µm from cell body center. H&I) Total dendritic length and number of branches per adult-born cell in GFP control and MC silenced mice (GFP control n = 11 cells/ 4 mice, MC silenced n = 15 cells/ 5 mice; *** p < 0.001, **** p < 0.0001). J) Sholl analysis demonstrates more extensive arborization of immature DGCs that developed in the absence of functional mossy cell inputs (*p= 0.03, 0.009, and 0.007 at intervals 20, 30, and 40µm distance from soma, **p= 0.007, 0.004, 0.001, 0.001, 0.001, 0.001, 0.004 at intervals 110, 120, 130, 140, 150, 160, and 170 µm distance from the soma, all other points not significant).

As an alternative approach, we used Cre-dependent viral vectors to silence hilar mossy cells with GFP-tagged tetanus toxin, and subsequently labeled adult born DGCs with a red fluorescent protein (mCherry)-expressing retrovirus. Comparable to findings after mossy cell ablation, we observed similarly enhanced dendritic outgrowth of 14-day old DGCs (Figure 2F-J). Again, cell proliferation and survival in the granule cell layer were not affected by mossy cell silencing (Supplemental Figure 4), but DCX+ cell density and outward migration demonstrated a mild but statistically significant increase in mice with permanently silenced mossy cells (Supplemental Figure 5D-F).

By 21 days post-mitosis, adult-born DGC dendrites reach the outer edge of the molecular layer and achieve a more mature branching pattern (van Praag et al., 2002; Zhao et al., 2006; Woods et al., 2018). Surprisingly, the dendritic morphology of 21-day old DGCs was not affected by either the absence of mossy cells or their functional silencing (Figure 3A-J). Together, these data indicate that mossy cell inputs primarily affect the maturation rate of adult-born granule cells, but not their mature dendritic pattern.

**Figure 3:**
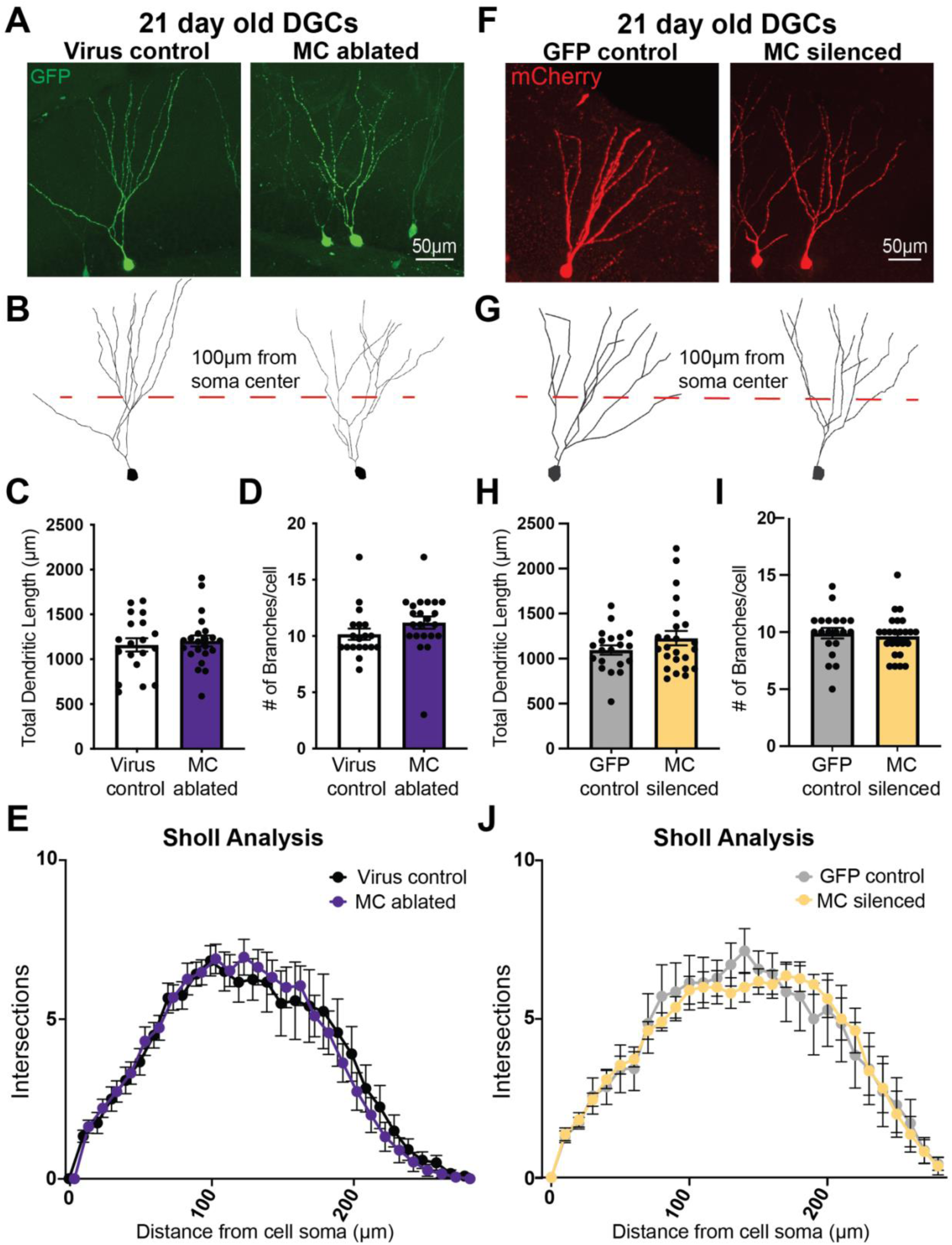
Adult-born DGCs achieve normal dendritic morphology in the absence of functional mossy cell inputs. A) Representative GFP-expressing adult-born dentate granule cells (DGCs) in virus injected control and mossy cell (MC) ablated mice, 21 days after retroviral labeling/mitosis. B) Skeleton tracing of cell morphology from the representative images in A, with a dashed line 100µm from cell body center. C&D) Total dendritic length and number of branches per adult-born cell in virus control and MC ablated mice (Virus control n = 19 cells/ 5 mice, MC ablated n = 22 cells/ 6 mice; p > 0.05). E) Sholl analysis demonstrates similar branching patterns of 21 day old DGCs in both groups. F) Representative mCherry-expressing adult-born (DGCs) in control (AAV5-flex-GFP) virus-injected and MC silenced (AAV5-flex-TeLC-injected) mice, 21 days after retroviral labeling. G) Skeleton tracing of cell morphology from the representative images in F, with a dashed line 100µm from cell body center. H&I) Total dendritic length and number of branches per cell of GFP control and MC silenced mice (GFP control n = 20 cells/ 6 mice, MC silenced n = 24 cells/ 6 mice; p >0.05). J) Sholl analysis demonstrates similar branching patterns of 21 day old DGCs in both groups.

### Dendritic spine formation in the IML persists in the absence of mossy cell inputs

Excitatory inputs in the molecular layer of the dentate gyrus originate from three distinct anatomical regions. Proximal IML inputs arise from hilar mossy cells, middle molecular layer (MML) inputs arise from medial entorhinal cortex, and distal outer molecular layer (OML) inputs arise from lateral entorhinal cortex. Spine densities on adult-born granule cell dendrites in each layer can provide insight into excitatory synaptogenesis (Woods et al., 2018). Dendritic spine formation is modulated by synaptic activity (McKinney et al., 1999; Toni et al., 2008; Kasai et al., 2010; Kim et al., 2018), and thus might be affected in the IML after mossy cell ablation or silencing.

Surprisingly, excitatory spine density of 21-day old DGCs in the IML was not altered following mossy cell ablation (Figure 4A,B). Similarly, spine density in the OML, from perforant path inputs, was also unchanged (Figure 4A,C). Given the extensive mossy cell ablation (Figure 1), the spine-like structures in the proximal molecular layer were either non-synaptic dendritic filopodia or associated with inputs from non-mossy-cells.

**Figure 4:**
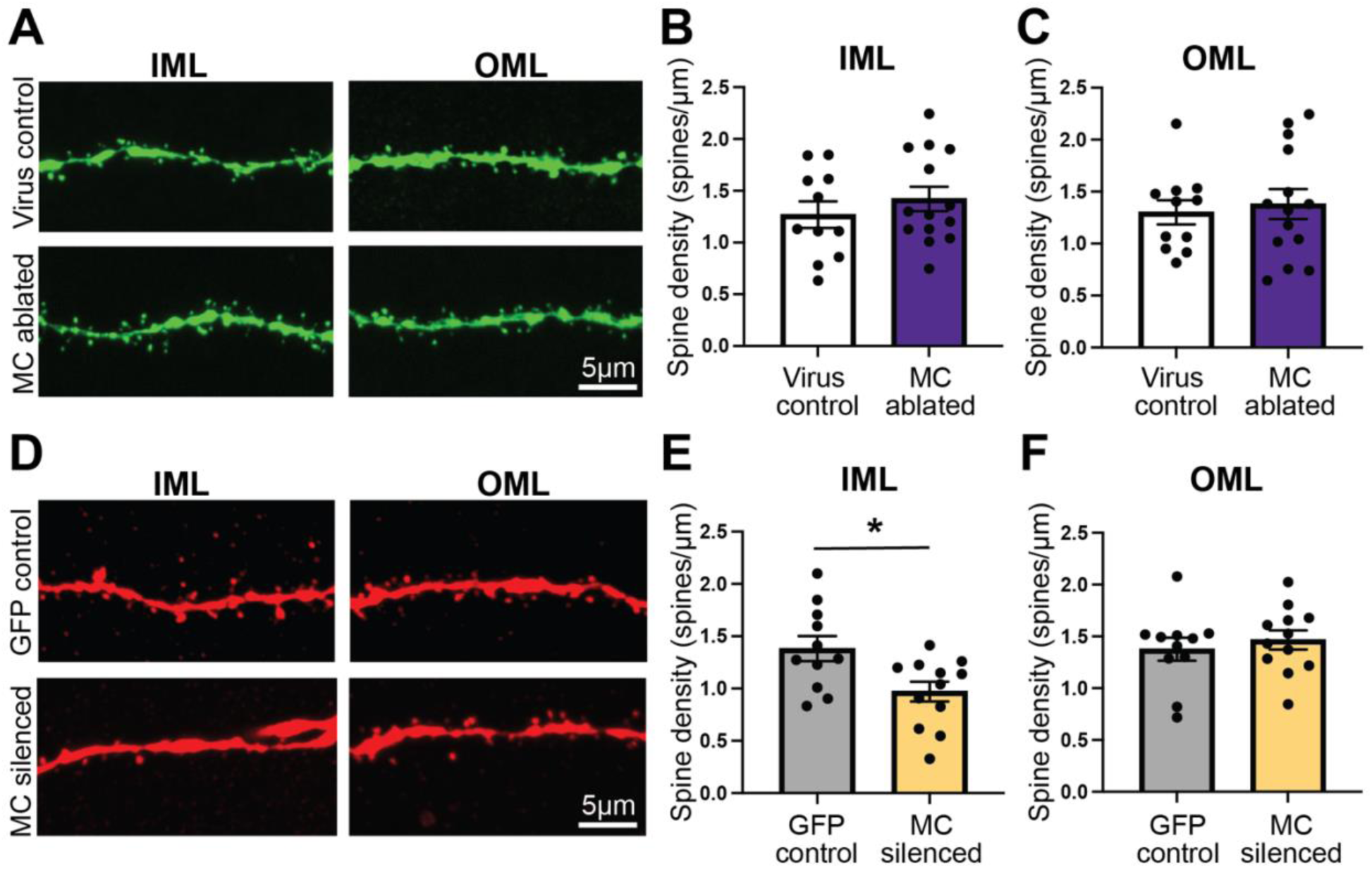
Differential effects of mossy cell ablation vs silencing on proximal dendritic spine formation by adult-born DGCs. A) Representative dendritic spines in the IML and OML from 21 day old, GFP-labeled adult born granule cells that developed in virus control and MC ablated conditions. B) IML spine density is not changed in cells born following mossy cell ablation (Virus control n = 11 cells/ 3 mice, MC ablated n = 14 cells/ 3 mice). C) OML spine density is not changed in cells born following mossy cell ablation (Virus control n = 11 cells/ 3 mice, MC ablated n = 14 cells/ 3 mice). D) Representative dendritic spines in the IML and OML from 21 day old, mCherry-labeled adult born granule cells that developed in GFP virus control and MC silenced conditions. E) IML spine density is reduced in cells born following mossy cell ablation (GFP control n = 11 cells/ 3mice, MC silenced n = 12 cells/ 3 mice; * p <0.05). F) OML spine density is not changed in cells born following mossy cell silencing (GFP control n = 11 cells/ 3mice, MC silenced n = 12 cells/ 3 mice).

In contrast, dendritic spine densities in 21-day old DGCs from mice with silenced mossy cells were reduced in the proximal molecular layer (Figure 4D,E), but not in OML dendrites (Figure 4D,F). The difference in spine density between these mossy cell manipulations could be indicative of differences in circuit modulation by the persistent yet silent mossy cell terminals.

### Functional integration of adult-born DGCs in the absence of mossy cell inputs

The integration of adult-born granule cells into the hippocampal network involves the stereotyped acquisition of excitatory and inhibitory synapses (Esposito et al., 2005; Ge et al., 2007; Zhao et al., 2008; Piatti et al., 2011). Although functional AMPA-receptor containing excitatory synapses parallel the development of dendritic spines, filopodial structures can lack pre-synaptic inputs (Toni et al., 2008; Piatti et al., 2011; Vivar et al., 2012; Sheng et al., 2018). We assessed excitatory synaptic function of retrovirally labeled (GFP or mCherry-expressing) 21 day-old adult-born granule cells and mature DGCs, 6 weeks after mossy cell ablation (van Praag et al., 2002), using whole-cell voltage clamp recordings of spontaneous excitatory post-synaptic currents (sEPSCs) and evoked synaptic responses in the IML. In mature DGCs, removal or silencing of mossy cell inputs reduced sEPSC frequency, but not sEPSC amplitude (Supplemental Figure 6), consistent with loss of excitatory synaptic innervation, similar to that previously observed (Jinde et al., 2012). Interestingly, however, sEPSCs in 3 week-old DGCs did not differ in frequency or amplitude from 3 week-old DGCs in age-matched control mice (Supplemental Figure 7). Thus, functional synaptic formation onto adult-born granule cells does not require mossy cell inputs.

Mossy cell axons drive direct excitation as well as feedforward inhibition of DGCs (Scharfman, 1995; Sloviter et al., 2003; Hsu et al., 2016; Butler et al., 2022). Despite the absence of mossy cell inputs, IML stimulation consistently evoked synaptic responses in both adult-born and mature DGCs (mean stimulus intensity: Virus control= 0.64 ±0.08 mA, MC ablated= 0.54 ±0.11 mA, MC silenced= 0.61 ±0.12mA; One way ANOVA, F(2,34)=0.2616, p=0.77). Surprisingly, the excitation:inhibition (E:I) ratio of IML-stimulated responses in 21 day-old DGCs from mossy cell ablated mice was elevated relative to control mice, suggesting a net loss of inhibitory synaptic drive (Figure 5C, D, G). In contrast, 21 day-old granule cells in mossy cell silenced mice had a E:I ratio similar to controls (Figure 5C, D, G). These data suggest that mossy cell axon loss alters the functional network integration of adult-born granule cells whereas mossy cell silencing did not. Evoked synaptic responses in mature DGCs in response to stimulation of the proximal IML also demonstrated an increase in E:I ratio from mossy cell ablated mice relative to mature DGCs in control mice (Figure 5A, B, E, F), whereas the E:I ratio of mature DGCs in mossy cell silenced mice was also unchanged, highlighting that these circuit changes are not exclusive to developing DGCs.

**Figure 5:**
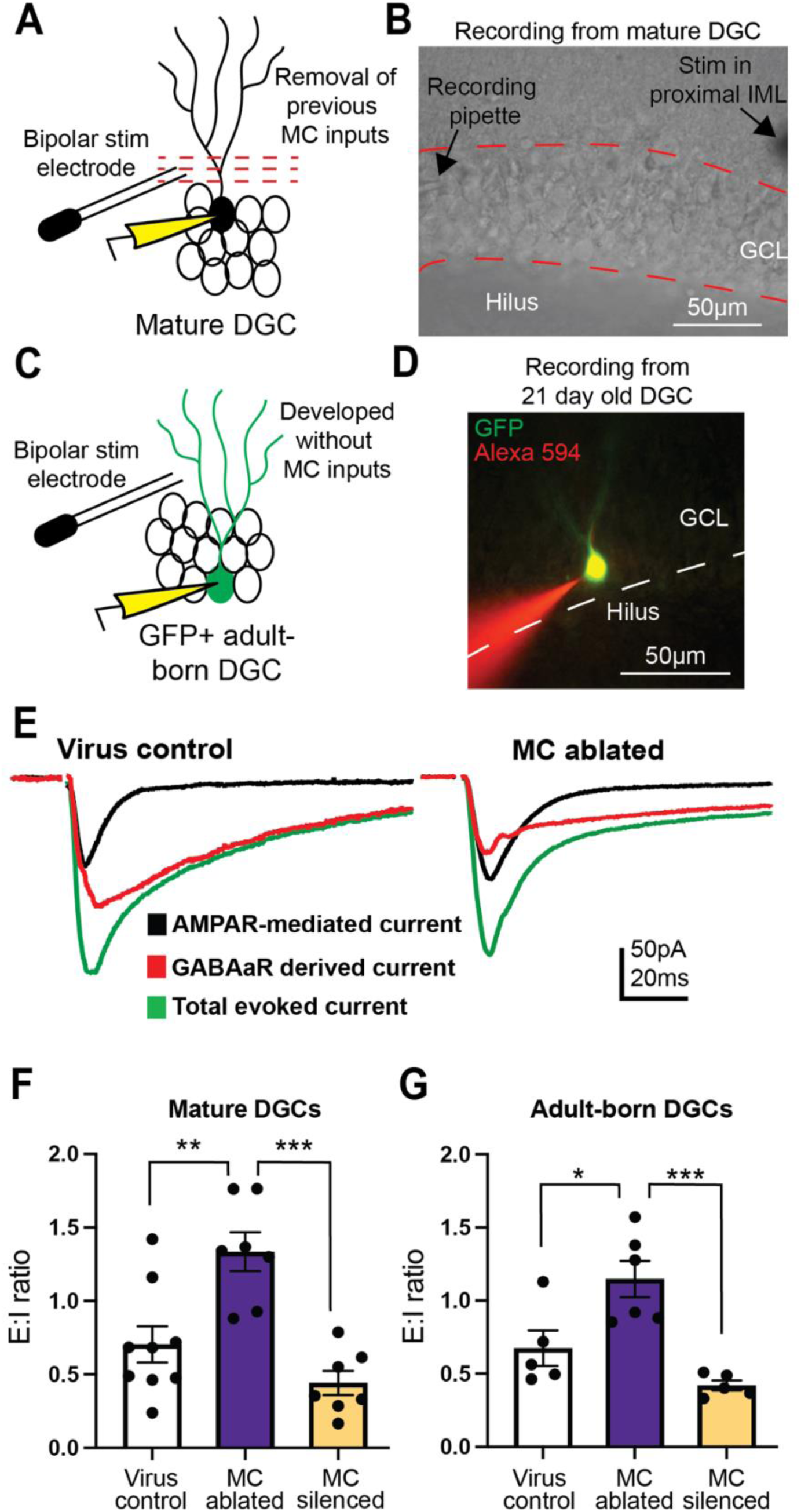
Altered evoked synaptic responses in both mature and adult-born DGCs following mossy cell ablation but not silencing. A) Schematic of recording paradigm for mature DGCs, with electrical stimulation of the proximal molecular layer. B) Representative image of the recording configuration for a mature DGC. C) Schematic of recording paradigm for virus-labeled 21 day old adult-born DGCs, with electrical stimulation of the proximal molecular layer. D) Representative image of a GFP+ 21dpi DGC during recording, filled with AlexaFluor 594 (red) to confirm cell specificity. E) Representative traces of total evoked synaptic current (green), AMPAR-mediated current (black), and GABAaR-mediated current (derived by subtraction, red) recorded from mature DGCs from virus control and mossy cell ablated conditions. F) Excitation:inhibition (E:I) ratios for synaptic responses to proximal molecular layer stimulation recorded from mature DGCs (Virus control n = 9 cells/ 7 mice, MC ablated n = 7 cells/ 6 mice, MC silenced n = 7 cells/ 6 mice; ** p<0.01, *** p<0.001). G) Excitation:inhibition (E:I) ratios of synaptic responses to proximal molecular layer stimulation recorded from 21 day old adult-born DGCs (Virus control n = 5 cells/ 3 mice, MC ablated n = 6 cells/ 3 mice, MC silenced n = 5 cells/ 5 mice; * p <0.05, *** p<0.001).

### Reorganization of the dentate gyrus molecular layer after mossy cell ablation

The persistence of evoked synaptic responses in the IML, together with the dendritic spines on newly born granule cells in the IML, suggests axonal reorganization. However, staining for the granule cell terminal marker zinc transporter 3 (ZnT3) did not reveal discrete ZnT3+ puncta in the IML of mice that had mossy cells ablated or permanently silenced at 6 weeks after virus injection (Supplemental Figure 8), or even later at 6+ months (data not shown), indicating that retrograde granule cell axon sprouting did not contribute to excitatory innervation after mossy cell ablation (Jinde et al., 2012).

In addition to cell body loss, mossy cell ablation caused dramatic thinning or even absence of mossy cell axon (calretinin) staining in the molecular layer (Figure 6A,B), and occurred across the dorsoventral axis of the hippocampus. This reduction in mossy cell axons was not observed 1 week after virus injection, but reduction in calretinin-positive axon staining was consistently observed 2, 3, and 6 weeks after virus injection suggesting that these axons are reduced early and unlikely undergoing further degeneration at later time points (Supplemental figure 9). In contrast, calretinin-labeled mossy cell axons were still present following mossy cell silencing, although the width of the calretinin-positive region of the molecular layer was slightly reduced and also contained enlarged axonal varicosities, consistent with the effects of tetanus toxin on synaptic vesicle accumulation (Figure 6A,B) (Woods et al., 2018). After ablation, any remaining mossy cell axons were tightly apposed to the granule cell layer, rather than more sparsely dispersed throughout the IML. This suggests that ablation either preferentially targeted cells projecting to the outer regions of the IML, or that the remaining mossy cell axons reorganized and migrated to more proximal locations.

**Figure 6:**
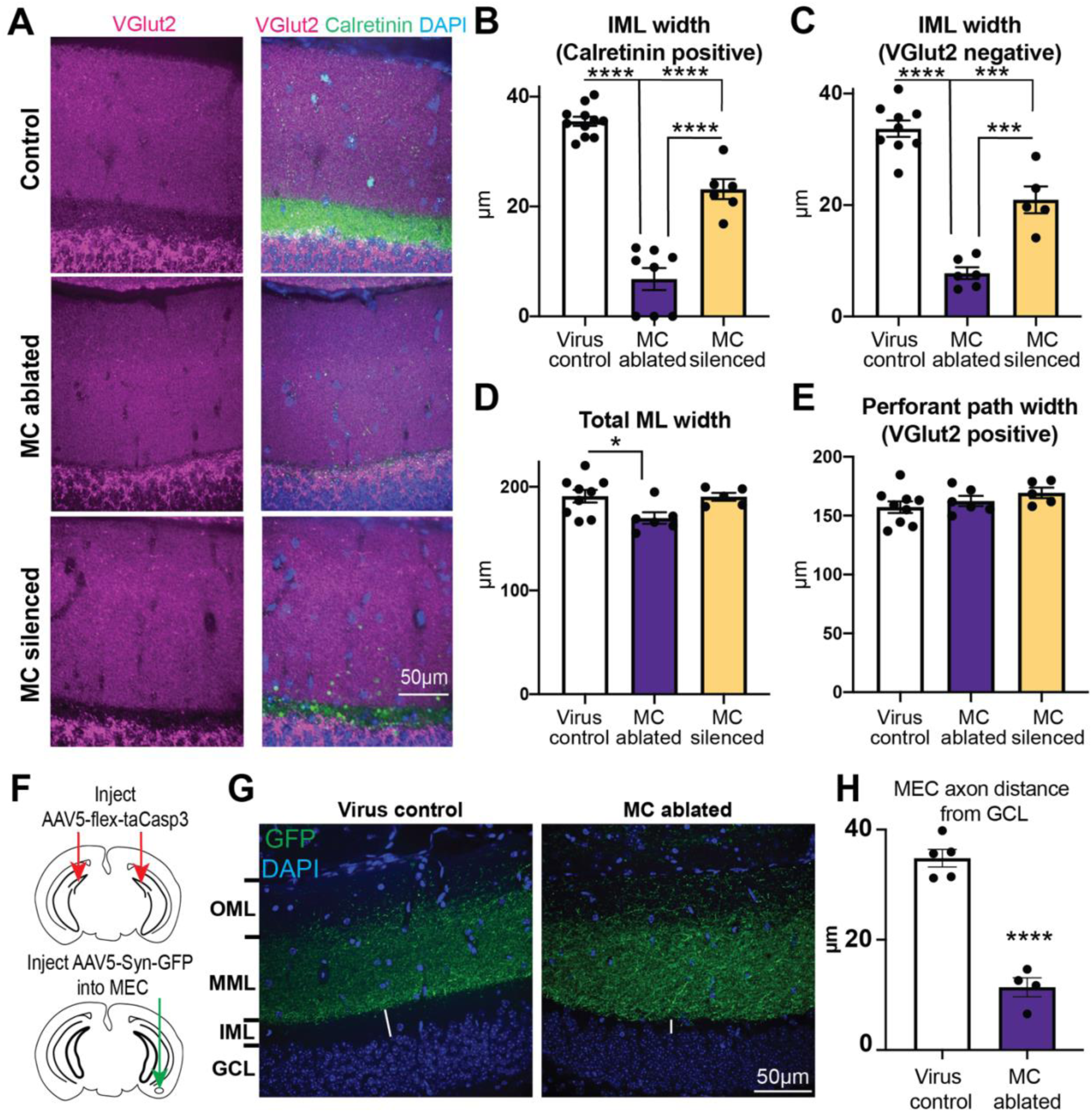
Mossy cell ablation causes collapse of the inner molecular layer. A) Representative images of VGlut2-expressing perforant path axons in relation to calretinin-expressing mossy cell axons from control, MC ablated, and MC silenced mice. B) IML width, as defined by calretinin staining (Virus control n = 9 mice, MC ablated n = 6 mice, MC silenced n = 5 mice, same mouse numbers for panels B-D; **** p < 0.0001). C) IML width, as defined by the VGlut2-negative region (*** p < 0.001, **** p < 0.0001). D) Total molecular layer (ML) width in control, MC ablated, or MC silenced mice (* p < 0.05). E) Perforant path width (VGlut2-positive middle and outer molecular layer) is unchanged after MC ablation, suggesting IML collapse. F) Schematic of viral injections to selectively label medial entorhinal cortex (MEC) axons in control and MC ablated mice. G) Representative GFP-positive MEC axons from virus control and MC ablated mice. White lines highlight the distance between labeled MEC axons and the granule cell layer (GCL) in control and MC ablated mice. H) Distance between the most proximal MEC axons and the GCL is reduced after MC ablation (Virus control n = 5 mice, MC ablated n = 4 mice; **** p < 0.0001).

To distinguish between these possibilities, we analyzed the location of perforant path synapses using anti-vesicular glutamate transporter 2 (VGlut2) immunohistochemistry, which selectively labels axon terminals deriving from the entorhinal cortex (Leranth and Hajszan, 2007; Perederiy et al., 2013). In control mice, dense VGlut2 staining was observed in the MML and OML, consistent with perforant path axon localization, with a clearly demarcated VGlut2-deficient region in the IML that coincided exactly with the location of calretinin-expressing mossy cell axons (Figure 6A). After mossy cell ablation, the width of the VGlut2-negative region of the molecular layer was dramatically reduced, and again nearly identical to the reduction in calretinin-positive layer thickness (Figure 6A,C). Mossy cell ablation also caused a small but statistically significant reduction in the total width of the molecular layer almost identical in absolute magnitude to the loss of IML width (Figure 6A,D). Thus, after ablation, perforant path synapses were now positioned adjacent to more proximal regions of the granule cell dendritic tree, as if the IML had collapsed.

Following mossy cell silencing, the VGlut2-negative region was only slightly reduced relative to virus controls, similar to the pattern observed for calretinin-positive axons in the proximal molecular layer (Figure 6A-C). The total molecular layer width was also unchanged in mossy cell silenced mice compared to virus control (Figure 6A,D), unlike mossy cell ablated mice. There was no significant difference in the width of the VGlut2-positive layers (perforant path) in any condition, indicating no compensatory expansion (Figure 6A,E). Together, these data demonstrate pronounced structural reorganization of the dentate gyrus molecular layer after mossy cell ablation, and more subtle effects after their permanent silencing.

VGlut2+ synapses in the proximal molecular layer after mossy cell ablation could represent *de novo* IML innervation by a non-entorhinal population of VGlut2-expressing terminals, or from translocation of MML inputs. To distinguish between these possibilities, we injected GFP-expressing viruses (either AAV5-CAG-GFP or AAV5-Syn-Chronos-GFP, in separate experiments) into the medial entorhinal cortex (MEC) in mice that had also received bilateral hippocampal injections of AAV5-flex-taCasp3 to ablate hilar mossy cells. In these mice, the location of the most proximal GFP-positive MEC axons was significantly closer to the molecular layer-granule cell border than control mice (Figure 6F-H). This result suggests that the more proximally located VGlut2+ synapses after mossy cell ablation reflected a shift in MEC fibers, again consistent with a “collapse” of the IML.

### Strongly reduced cannabinoid sensitivity of proximal excitatory synapses after mossy cell removal

Mossy cell axon terminals express presynaptic cannabinoid receptors (CB1Rs), which potently suppress neurotransmission from mossy cells onto target granule cells (Katona et al., 2006; Monory et al., 2006; Uchigashima et al., 2011; Jensen et al., 2021), whereas perforant path synapses do not express CB1Rs (Monory et al., 2006). As expected (Chiu and Castillo, 2008), bath application of WIN 55,212-2 strongly suppressed EPSC amplitudes evoked after IML stimulation in mature DGCs from control mice (Figure 7). In contrast, the WIN sensitivity of proximally-evoked EPSCs in mature DGCs from either mossy cell ablated or silenced mice was substantially reduced (Figure 7A-C), consistent with the loss or silencing of mossy cell inputs. This relationship was also observed in adult-born DGCs (Figure 7D), consistent with functional innervation of the IML by WIN-insensitive inputs. The small residual CB1R-sensitivity observed likely represents a small population of non-ablated/non-silenced mossy cell fibers. These data indicate functional and structural reorganization of the IML after mossy cell ablation, with preserved innervation of granule cell dendrites in the IML by axons from the entorhinal cortex.

**Figure 7:**
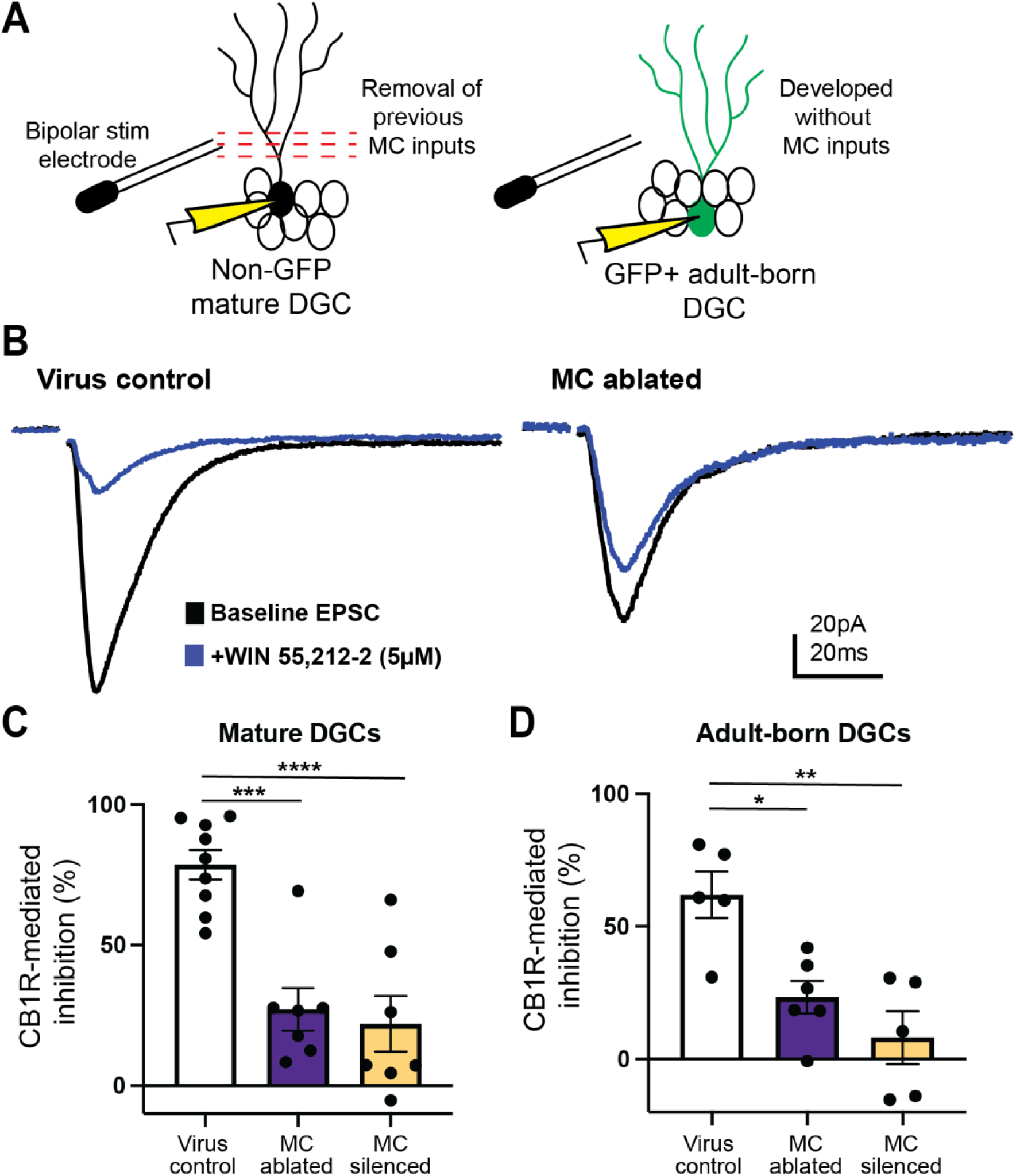
Strongly reduced CB1R-mediated inhibition of proximal molecular layer inputs after mossy cell ablation or silencing. A) Schematic of recording paradigm. A stimulating electrode was used to activate inputs in the proximal molecular layer, while recording from mature or retrovirally-labeled adult born DGCs from control, MC ablated or MC silenced mice. B) Representative AMPAR-mediated evoked excitatory synaptic responses before and after application of WIN 55,212-2 (5µM). C) EPSCs recorded from mature DGCs in both MC ablated and MC silenced mice had minimal WIN sensitivity relative to cells from control mice (Virus control n = 9 cells/ 7 mice, MC ablated n = 7 cells/ 6 mice, MC silenced n = 7 cells/ 6 mice; *** p <0.001, **** p <0.0001). D) EPSCs recorded from 21 day old DGCs in MC ablated and MC silenced mice had minimal WIN sensitivity relative to control cells (Virus control n = 5 cells/ 3 mice, MC ablated n = 6 cells/ 3 mice, MC silenced n = 5 cells/ 5 mice; *p < 0.05; ** p < 0.01).

### Mossy cell loss or silencing does not affect granule cell activity or seizure susceptibility

Hilar mossy cells can modulate hippocampal activity through long-range projections (Buckmaster and Schwartzkroin, 1994; Scharfman, 1995; Buckmaster et al., 1996). To determine whether mossy cell loss or silencing globally alters dentate gyrus activity, we stained for the activity-dependent immediate early gene product, cFos (Morgan et al., 1987; Harvey and Sloviter, 2005). There was no difference in basal cFos expression between experimental groups (Figure 8), suggesting that mossy cells either do not modulate basal granule cell activity, or that the dentate gyrus compensated for the acute effects of mossy cell loss or silencing.

**Figure 8:**
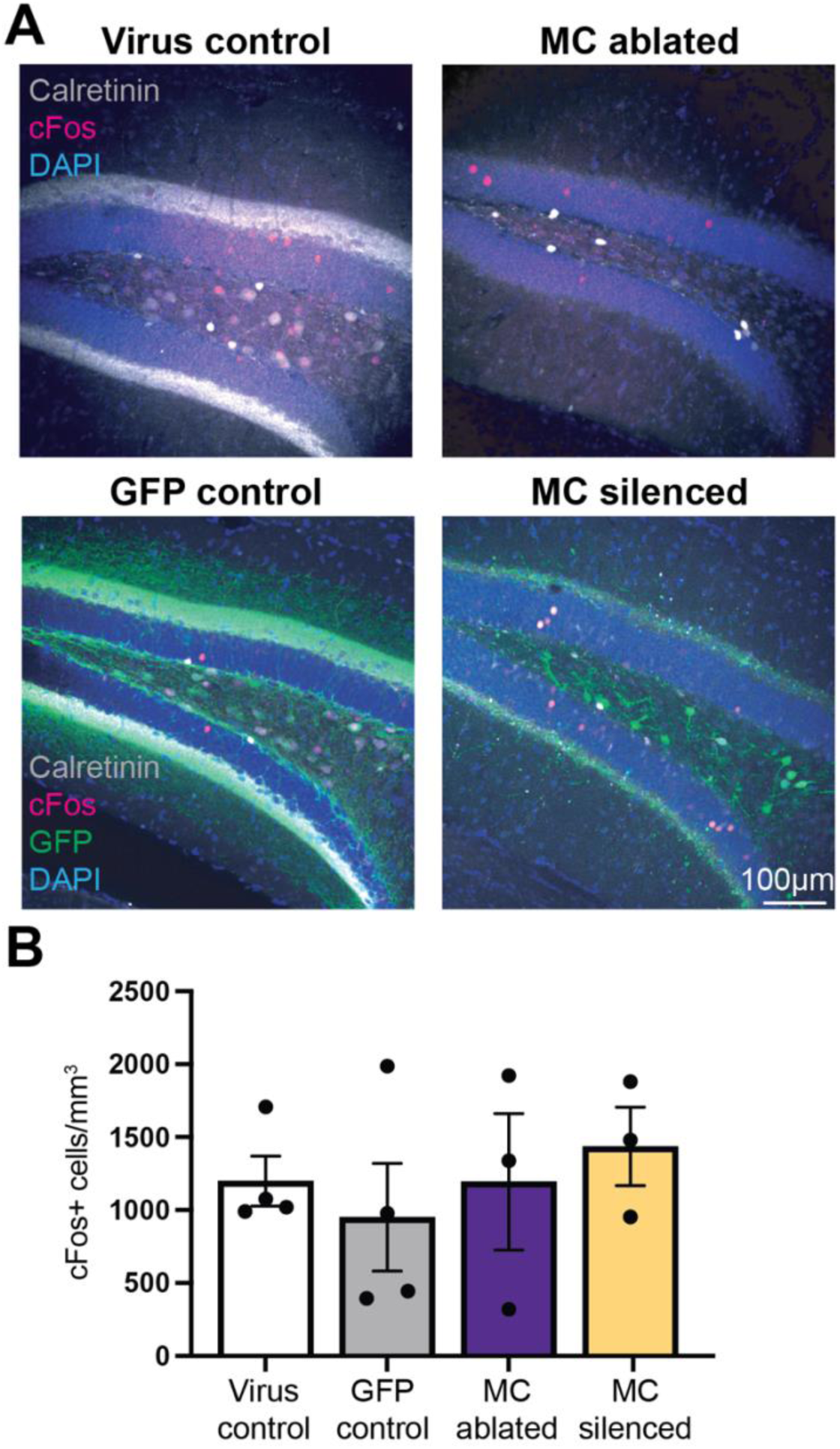
Basal dentate granule cell activity is not altered by functional mossy cell loss. A) Representative cFos expression (red) in dentate granule cells in virus injected Cre-negative control, GFP virus injected control, MC ablated or MC silenced mice. Calretinin staining (white) of the IML highlights mossy cell axons, while calretinin staining highlights immature neuroblasts in the subgranular zone and mossy cell somata in the hilus, respectively. B) cFos-positive cell density in the dentate granule cell layer; (Virus control n = 4 mice, GFP control n = 4 mice, MC ablated n = 3 mice, MC silenced n = 3 mice; n.s. all comparisons).

Mossy cells acutely control seizure initiation and duration in mouse models of seizure susceptibility or epilepsy (Jinde et al., 2012; Bui et al., 2018; Botterill et al., 2019). Additionally, the elevated E:I ratio at IML inputs after mossy cell ablation (Figure 5) could suggest reduced inhibitory tone. To determine whether this was associated with a reduced seizure threshold, we administered the convulsant pentylenetetrazol (PTZ, 40mg/kg i.p.) to mice 6 weeks after control virus injections, mossy cell ablation, or mossy cell silencing, with a subset of mice receiving only saline injections (i.p.) as an additional control.

PTZ caused behavioral seizures in all groups, and consistently activated DGCs as assessed by cFos staining (Figure 9A,B). However, there was no difference in maximum seizure scores in any group of PTZ-treated mice (Figure 9C), all of which were significantly greater than saline-treated controls. There also was no difference in cumulative seizure score or in the time to first behavioral seizure after PTZ injection between mice with intact mossy cells when compared with mice after mossy cell ablation or silencing (Figure 9D,E), indicating that chronic mossy cell loss or silencing did not alter induced seizure susceptibility. This is consistent with observations after *chronic* mossy cell removal, and notably different from the increased seizure susceptibility caused by *acute* mossy cell inactivation (Jinde et al., 2012; Bui et al., 2018; Botterill et al., 2019).

**Figure 9:**
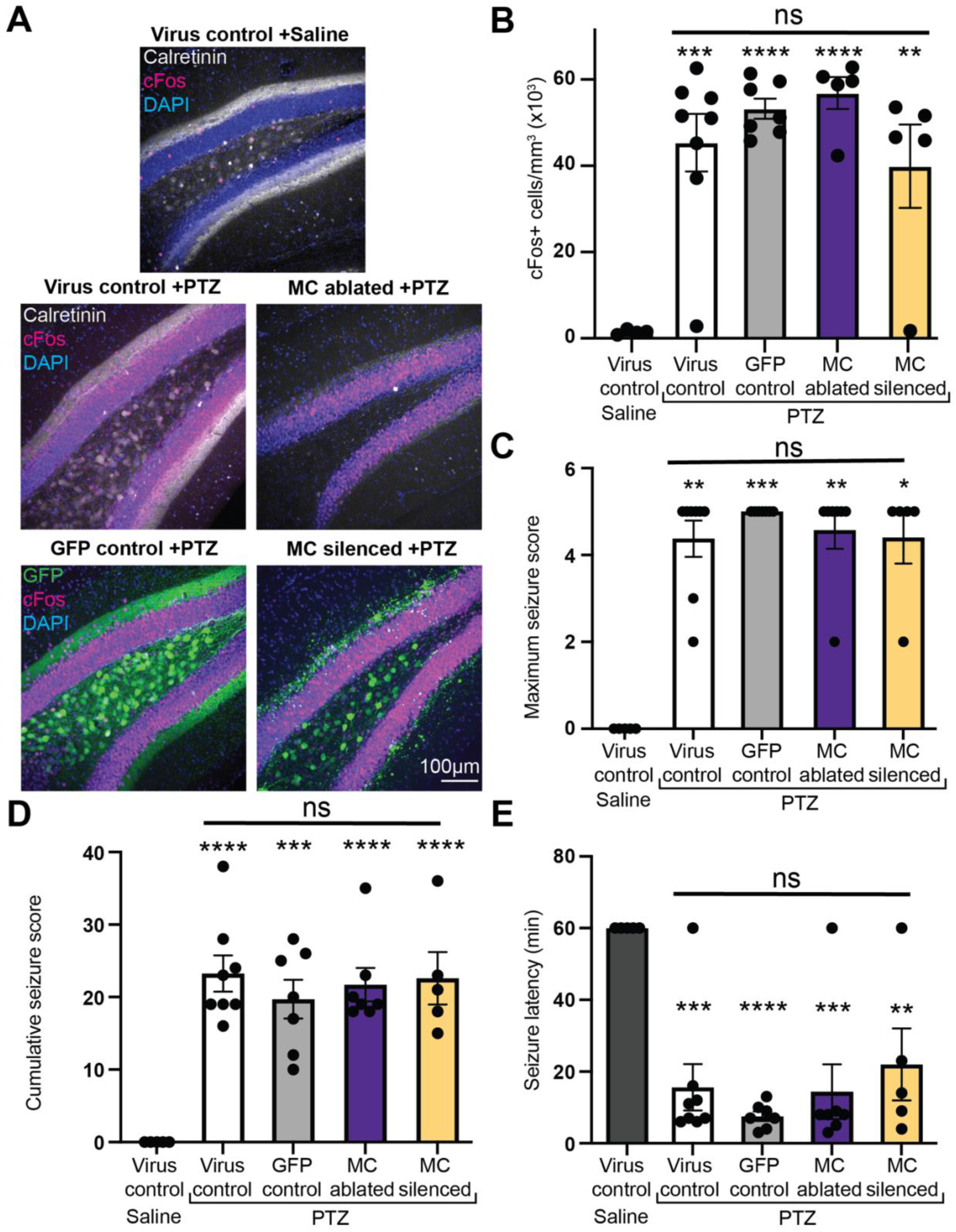
Neither mossy cell loss nor silencing alters seizure susceptibility. A) Representative cFos expression (red) in dentate granule cells after saline or PTZ injection. B) Granule cell cFos expression is increased after PTZ injection similarly in all groups (Virus control Saline= 5 mice, Virus control PTZ= 8 mice, GFP control PTZ= 7 mice, MC ablated PTZ= 7 mice, and MC silenced PTZ= 5 mice; ** p <0.01, *** p <0.001, and **** p <0.0001 compared to saline; n.s. for all PTZ-treated groups relative to each other). C) Maximum seizure scores (* p<0.05, ** p <0.01, and *** p <0.001 compared to saline; n.s. for all PTZ-treated groups relative to each other). D) Cumulative seizure scores (*** p <0.001, and **** p <0.0001 compared to saline; n.s. for all PTZ-treated groups relative to each other). E) Latency to first seizure (time to S3 or greater on modified Racine scale) (** p <0.01, *** p <0.001, and **** p <0.0001 compared to saline; n.s. for all PTZ-treated groups relative to each other).

## Discussion

Hilar mossy cells are increasingly recognized for their roles controlling hippocampal function and behavior (Bui et al., 2018; Botterill et al., 2019; Fredes and Shigemoto, 2021; Jensen et al., 2021; Noguchi et al., 2023; Alcantara-Gonzalez et al., 2025). By selectively removing or silencing mossy cells we identified important and unexpected roles of mossy cells in maintaining hippocampal circuit organization, while also demonstrating the ability of the hippocampus to compensate for mossy cell loss.

### Reorganization of the dentate molecular layer in the absence of mossy cell inputs

Mossy cell loss dramatically altered the anatomical and functional organization of the dentate molecular layer, collapsing more distal entorhinal inputs to more proximal regions, without causing any compensatory granule cell axon sprouting. This “IML collapse” following hilar mossy cell ablation was novel and unexpected. Although hilar mossy cell loss occurs in models of acquired epilepsy and TBI, subsequent excitatory axonal sprouting or reorganization occurs in the same layer as lost mossy cell axons (Lowenstein et al., 1992; Buckmaster and Dudek, 1997; Sloviter et al., 2003; Overstreet-Wadiche et al., 2006). It is unclear whether the more limited mossy cell loss in brain injuries (∼50%) prevents IML collapse, or whether reinnervation by sprouted granule cell axons preserves this layer after injury. In the setting of partial (∼35%) mossy cell input loss due to contralateral hippocampal removal, IML thickness is preserved due to reinnervation by the surviving ipsilateral mossy cells (McWilliams and Lynch, 1978), suggesting that IML preservation after injury could involve both mechanisms.

Molecular layer lamina in the dentate gyrus appear to have differential capacities for re-organization in the context of injury or neuronal disease. Lesions to the perforant path cause extensive dendritic spine loss by mature DGCs, even though the granule cells are otherwise preserved and eventually re-innervated, possibly via commissural/mossy cell axons (Matthews et al., 1976b, a; Steward et al., 1988; Deller and Frotscher, 1997). Interestingly, selective lesion of the MEC/MML actually drives widening of the IML (Phinney et al., 2004), and layer-specific IML widening can also occur in a mouse model of Alzheimer’s disease (Alcantara-Gonzalez et al., 2025) associated with loss of perforant path synapses (Stranahan and Mattson, 2010). Taken in context with our current data, it is clear that modes of plasticity and capacities for reorganization likely reflect intrinsic differences between layers as well as different responses to distinct injuries and pathologies.

Although molecular mechanisms underlying layer-specific reorganization during both development and circuit rearrangement have yet to be identified, target and input-specific expression of synaptic adhesion molecules are prime candidates for controlling this specificity (Jucker et al., 1996; Fasen et al., 2002). The dendrites of granule cells that acquire more proximal MEC inputs following mossy cell ablation are functionally distinct given the lack of WIN-sensitivity and elevated E:I ratios at excitatory synapses in that region. Thus, one can conclude that cell-autonomous, input-specific factors (such as expression of specific adhesion molecules and receptors) critically contribute to functional innervation, and that different layers have layer-specific cues that guide both organization and re-innervation (Stranahan and Mattson, 2010).

### Role of mossy cells in adult-born DGCs

The transient acceleration of dendritic outgrowth from adult-born DGCs after removal or silencing of hilar mossy cells highlights the fundamental influence that mossy cells provide to adult-born DGCs during their early dendritic development, and supports the hypothesis that aberrant outgrowth of immature DGCs observed in translational models of brain injury is in part driven by mossy cell dysfunction (Overstreet-Wadiche et al., 2006; Niv et al., 2012; Butler et al., 2015; Villasana et al., 2015; Tensaouti et al., 2020). Interestingly, the transient acceleration of dendritic outgrowth is opposite of the effect of perforant path lesions, which reduce dendritic outgrowth of adult-born DGCs (Perederiy et al., 2013). This difference could be explained by a mossy cell-derived signal that stalls dendritic development in the proximal molecular layer before subsequent outgrowth into more distal regions of the molecular layer. Although it is unknown what this signal might be, the persistence of accelerated dendritic outgrowth in the presence of silenced mossy cells - in which presynaptic terminals remain present - suggests that a secreted signal (such as glutamate) might critically control the rate of dendritic outgrowth. Given the relative importance of perforant path inputs to the later stages of adult-born granule cell integration (Schmidt-Hieber et al., 2004; Vivar et al., 2012; Woods et al., 2018), it is perhaps not surprising that the persistent perforant path inputs stabilize normal patterns of DGC dendritic growth.

Interestingly, the proximal dendritic spine density of adult-born DGCs was unchanged following mossy cell ablation but reduced when mossy cells were permanently silenced. Given the “collapse” of the former IML after mossy cell ablation, it is clear that mossy cell axons are not specifically required for synapse formation onto proximal inputs, as perforant path axons were able to compensate for the lost fibers in this region. Additionally, the reduced spine density in the presence of silenced mossy cell axons suggests that activity-dependent mechanisms contribute meaningfully to spine formation or maintenance. It is unclear whether the remaining IML spines after silencing represent filopodia-like structures without associated presynaptic terminals, or the connection of spine structures with silenced terminals, similar to observations made after perforant path silencing (Woods et al., 2018). In both cases, however, sEPSC frequencies onto recently developed DGCs were unchanged, suggesting that the total functional innervation of each newly generated granule cell might be controlled during their maturation in a cell-autonomous and homeostatic manner. Interestingly, unlike the compensatory increase in IML spine density and shift towards larger and faster events seen after perforant path lesions (Perederiy et al., 2013), distal perforant path spine density was unchanged after IML lesion, suggesting different regulatory mechanisms.

### Acute vs chronic mossy cell impairment

Classically, there have been two main hypotheses regarding the role of mossy cells in seizures and epilepsy: the irritable mossy cell hypothesis and dormant basket cell hypothesis (Sloviter, 1987; Santhakumar et al., 2000; Sloviter et al., 2003). In the irritable mossy cell hypothesis, residual mossy cells become hyperactive after an initial insult and contribute to seizures by driving granule cell activation. Alternatively, by focusing on the important role that mossy cells play in driving inhibitory basket cell function, mossy cell loss after seizures could cause basket cells to become “dormant”, thus disinhibiting the circuit and making subsequent seizures more likely.

More recent work, however, has suggested that the involvement of mossy cells during seizures can depend on the stage of epileptogenesis (Jinde et al., 2012; Bui et al., 2018; Botterill et al., 2019; Butler et al., 2022). For example, when mossy cells are chemogenetically inhibited in an otherwise intact/healthy circuit, the induction of status epilepticus in mice by pilocarpine is inhibited, which then reduces the incidence and severity of subsequent seizures (Botterill et al., 2019). This suggests that mossy cell activity might acutely contribute to induced seizures in otherwise healthy mice, perhaps through its recurrent excitatory connections with granule cells. Although this was not observed when mossy cells were acutely depleted in *ex vivo* hippocampal slices (Ratzliff et al., 2004), differences in the experimental approaches (intact vs reduced circuits) could explain this apparent discrepancy.

In contrast, mossy cells that survive pilocarpine-induced status epilepticus have enhanced excitatory connectivity to parvalbumin-positive basket cells (Butler et al., 2022), which may explain why optogenetic stimulation of surviving mossy cells in epileptic mice suppresses seizure frequency (Bui et al., 2018). This apparently contradictory finding may represent circuit rearrangements which develop during epileptogenesis. Consistent with this concept, when mossy cells were ablated from mice using a diphtheria-toxin-mediated strategy, seizure susceptibility to kainic acid was transiently elevated (4-11 days after ablation) but returned to baseline 6-8 weeks after ablation (Jinde et al., 2012). Although the transiently increased seizure susceptibility could alternatively be explained by the expression of kainate receptors by residual/dying mossy cells (Ramos et al., 2022), altered mossy cell involvement during the acute and chronic phases of epileptogenesis may explain the apparent discrepancies.

Although the imbalance of synaptic excitation and inhibition is hypothesized as a root cause of neurological and psychiatric disease (Ziburkus et al., 2013), its broader impact on circuit function can be difficult to predict. We observed altered synaptic E:I ratios after mossy cell ablation, but this did not translate to broader increases in granule cell activity. Possible reasons for this discrepancy could be that mossy cell-driven feedforward inhibition is not necessary for general inhibitory tone in the hippocampus, or that non-synaptic mechanisms such as increased tonic inhibition could provide the necessary functional compensation (Boychuk et al., 2016; Parga Becerra et al., 2021; Boychuk et al., 2022). Together, these observations suggest the dentate circuit can functionally and structurally reorganize in the face of changes which acutely alter circuit excitability, with the remaining circuits able to achieve a new homeostatic balance in which granule cell activity remains low. However, when coupled with additional injury-induced circuit rearrangements, the circuit may no longer effectively compensate, in which case epileptogenesis could occur. Identification of the mechanisms involved in both the restoration of this homeostatic balance as well as the injury-associated signals that prevent compensation will hopefully lead to better strategies to reduce seizures and the development of epilepsy.

## Materials and Methods

### Animals

Transgenic mice selectively expressing Cre in hilar mossy cells under the control of the Calcitonin receptor-like receptor (Crlr) promoter were created as previously described (Jinde et al., 2012) and obtained from Jackson Laboratory (Bar Harbor, ME; C57BL/6N-Tg(Calcrl,cre)^4688Nkza/J^; stock # 023014). Crlr-Cre mice were maintained on a C57BL/6NJ background with intermittent backcrossing to wildtype C57BL/6NJ mice. Mice were housed with a 12 hr light/12 hr dark cycle in the Oregon Health & Science University or Portland VA vivarium (both AAALAC-approved), with food and water provided *ad libitum.* Both male and female mice were used for experiments. All procedures were approved by the Oregon Health & Science University and Portland VA Institutional Animal Care and Use Committees and followed NIH guidelines for the care and treatment of animals.

### Viral Vector Delivery

As we observed Cre-mediated recombination in distal CA3 pyramidal neurons and Purkinje cells of the cerebellum after crossing Crlr-Cre mice with tdTomato reporter mice (unpublished observation), we used stereotaxic dentate hilus injections of Cre-dependent viral vectors to minimize off-target expression of Cre-dependent proteins (Butler et al., 2022). Briefly, mice were anesthetized using 4% isoflurane, after which anesthesia was maintained with inhaled 1-2% isoflurane and continuous monitoring. After anesthetic induction, the mouse’s head was shaved and mounted onto the stereotaxic frame, after which a sterile midline scalp incision was made, followed by stereotaxic drilling. To target hilar mossy cells, mice were bilaterally injected with 1 µL AAV-containing saline delivered via Hamilton syringe at 0.25µL/min, using coordinates measured from bregma: ±2.35mm X, -2.80mm Y, and -2.70mm Z. To target medial entorhinal cortex pyramidal neurons, a subset of mice were injected with 1µL of non-Cre dependent AAV vectors with a GFP fluorescent tag at 0.25µL/min to label axons projecting to the dentate gyrus, with coordinates measured from lambda: ±3.4mm X, - 0.7mm Y, -2.5 mm. For retroviral labeling of adult-born granule cells, coordinates were measured from bregma: ±1.1mm X, -1.9mm Y, -2.5 mm & -2.3mm Z, with a volume of 1μL injected at each site at 0.25μL/min (2 injections per hemisphere, targeting both blades of each dentate).

After each injection, the needle remained in place for at least one minute, after which it was slowly removed. The scalp incision was closed with surgical glue (Vetbond). Mice were given soft food with 1:10 diluted flavored Tylenol solution (3.2 mg/mL) to alleviate pain following surgery, and mice were monitored every 24hrs for 3-4 days after injection.

### Mossy cell ablation and silencing

To selectively ablate mossy cells in Crlr-Cre mice, we bilaterally injected Cre-dependent AAV-derived viral particles expressing activated caspase-3 (AAV5-flex-taCasp3-TEVp; 1 x 1012 particles/ml; UNC viral core, Chapel Hill, NC (Yang et al., 2013); Figure 1A). These vectors encode a designer form of caspase 3 that contains the cleavage site for tobacco etch virus protease (TEVp) fused to pro-caspase 3, together with the TEVp protease in a bicistronic arrangement; infected Cre-expressing cells express TEVp, which cleaves pro-caspase3 into taCasp3 and activates cell death via apoptosis (Yang et al., 2013). As controls, we injected this same virus in equivalent amounts/coordinates in age-matched WT (i.e., Crlr-Cre-negative) mice. For a subset of mice injected with AAV5-flex-taCasp3-TEVP virus into the dentate hilus, a subsequent injection, during the same surgery but with additional craniotomy sites, of either AAV5-Syn-Chronos-GFP (UNC viral core) or AAV5-CAG-GFP (Addgene, Watertown, MA, plasmid #37825) was delivered into the medial entorhinal cortex to label the axons from these neurons.

In separate experiments, we permanently silenced mossy cells in Crlr-Cre mice by injecting a Cre-dependent virus expressing tetanus toxin light chain (TeLC) protein fused to GFP (AAV5-flex-TeLC-GFP; 1 x 1013 particles/ml). A Cre-dependent virus expressing green fluorescent protein (GFP) was used as a control (AAV5-flex-GFP; 1 x 1013 particles/ml). The plasmids for AAV5-flex-GFP and AAV5-flex-TeLC-GFP were generously provided by Dr. Peer Wulff (Murray et al., 2011) and packaged by VectorBuilder (Chicago, IL).

### Retroviral vector production and labeling of adult-born granule cells

Moloney Murine Leukemia Virus (MMLV)-based retroviral vectors selectively infect mitotic cells, and are an established technique for labeling adult-born DGCs (van Praag et al., 2002). GFP- or mCherry-expressing retroviral particles were created using pRubi-based vectors and transfection of GP2 cells as previously described (Luikart et al., 2012), and concentrated to ∼1 x 107 particles/ml using RetroX Concentrator (Takara Bio, Clontech). Retroviral injections occurred 3 weeks after Cre-dependent AAV or control virus injections.

### BrdU labeling of mitotic cells

To label mitotic cells, we administered BrdU (300mg/kg i.p. for each of 2 injections 4 hours apart) to a subset of mice 3 weeks after intrahippocampal AAV injection. These mice were perfusion-fixed at 24hr or 21 days following BrdU injection, to assess proliferation or survival of adult-born DGCs, respectively.

### Perfusion fixation

Perfusion fixation was performed on terminally anesthetized mice after inhaled isoflurane anesthesia followed by i.p. injection of 2% Avertin (0.8mL). Mice were then transcardially perfused with 10 mL ice-cold 0.1M phosphate-buffered saline (PBS) followed by 15 mL fixative (4% paraformaldehyde in 0.1M PBS, pH ∼7.3-7.4). After dissection, brain tissue was post-fixed overnight at 4oC followed by PBS rinsing. Free-floating coronal brain sections were made using a Leica VT1000 vibratome at 100µm thickness except for retrovirus-injected mice, which were sectioned at 150µm to preserve dendritic trees.

### Immunohistochemistry

To assess mossy cell ablation, axonal reorganization, and glial activation, free-floating brain sections (4 sections 600µm apart for 100µm thick sections and 3 sections 900 µm apart for 150 µm thick sections) were permeabilized and blocked using 0.1M PBS with 0.4% Triton-X 100 (PBST) containing 10% normal horse or goat serum for 1hr at room temperature. Following permeabilization, tissue was stained overnight at 4oC in PBST + 1.5% normal horse serum and primary antibodies. Primary antibodies used included: anti-calretinin (1:1000, goat, Swant, CG1), anti-GluA2 (1:500, rat, Synaptic Systems 182 117), anti-ZnT3 (1:500, guinea pig, Synaptic Systems, 197 004), anti-VGlut2 (1:500, guinea pig, Synaptic Systems, 135 418), anti-cFos (1:500, guinea pig, Synaptic Systems, 226 308), anti-Iba1 (1:500, rabbit, Fujifilm Wako, 016-26721), anti-galectin 3 (1:500, goat, Novus Biological, MAB1197), anti-DsRed (1:500, rabbit, Takara Bio 632496) and anti-GFP (1:1000, chicken, Aves Labs, GFP-1020, or Alexa Fluor 488-conjugated anti-GFP, 1:500, rabbit, Invitrogen A-21311). Sections were then rinsed with PBST 3x 5min and incubated for 4-6hr at room temperature in PBST with 1.5% normal horse serum and secondary antibodies: 1:400 donkey anti-rabbit Alexa 647 (Invitrogen, A31573), 1:400 donkey anti-chicken Alexa 488 (Jackson ImmunoResearch Laboratories, 703-548-155), 1:400 donkey anti-goat Alexa 647 (Invitrogen, A21447), donkey anti-rabbit Alexa 488 (Invitrogen, A21206), goat anti-rat 647 (Jackson ImmunoResearch Laboratories, 112-605-003),and/or donkey anti-guinea pig Cy3 (Jackson ImmunoResearch Laboratories, 706-166-148). After staining, sections were incubated in 1:20K DAPI for nuclear staining, then mounted onto Fisher Superfrost slides with Fluoromount G (Southern Biotech, 0100-01).

To assess cell proliferation and neurogenesis, we used free floating sections (4 sections 600µm apart) permeabilized with 0.1M KPBS with 0.4% Triton X-100 (KPBST, pH=7.4) at room temperature for 45 minutes prior to a 30 minute incubation in KPBST with 2N HCl at 37oC, followed by wash in KPBST pH=8.5 at room temperature for 10 min and two washes in KPBST pH=7.4 for 10 minutes at room temperature. Tissue was then incubated with KPBST containing 10% horse serum for ∼1hr and then placed in staining solution (KPBST+1.5% horse serum) with primary antibodies against doublecortin (1:500, guinea pig, Millipore ab2253) and BrdU (1:500, rat, Abcam ab6326) overnight at 4oC. Sections were then rinsed with KPBST 3x 5min and incubated for 4-6hr at room temperature in KPBST with 1.5% normal horse serum and secondary antibodies: 1:400 donkey anti-rat Alexa 647 and 1:400 donkey anti-guinea pig Alexa 568. Tissue was rinsed, counterstained with DAPI and mounted onto Fisher Superfrost slides with Fluoromount G (Southern Biotech).

### Confocal microscopy and image analysis

Images were acquired using a Zeiss LSM 780 or 900 laser scanning confocal microscope on a motorized Axio Observer Z1 inverted scope (Carl Zeiss MicroImaging) or a Yokogawa CSU-W1 spinning disk confocal microscope mounted on a Scientifica Frame with Olympus optics (Intelligent Imaging, Inc). Images were taken with 20X 0.8NA (air) or 40X 1.4NA (oil) objectives. For cell density measurements (mossy cell, doublecortin, BrdU, and cFos), a 20X confocal stack (20-30µm depth; 0.63µm step) was acquired for analysis from 3-4 sections per animal, the observations made in these individual sections were then averaged to generate a single value for each mouse (n = # of mice). For CA3 cell counts in mossy cell ablated mice, images of DAPI-stained CA3 were taken with a 40X 1.4NA (oil) objective, and a single section plane (3-4 sections per animal) was imaged in the proximal portion of the CA3 pyramidal cell layer (∼100µm away from the hilus). For GFP+ cell counts of CA3 neurons in GFP control and mossy cell silenced mice, images were taken with a 20X 0.8NA (air) objective and a single section plane (3-4 sections per animal) was imaged in proximal CA3. Cell counts were manually performed in FIJI by a blinded observer using the Cell Counter plugin in FIJI (National Institutes of Health) and divided by the granule cell layer volume imaged for each section.

To assess mossy cell density after viral injection, we measured calretinin-positive cell bodies in the hilus of the 2 most ventral sections of the 4 sections of stained tissue, coinciding with regions of highest calretinin expression (Houser et al., 2020; Botterill et al., 2021). Additionally, in a subset of these mice, we measured GluA2 hilar cell density in the two most dorsal hippocampal tissue sections to assess if our virus-mediated mossy cell ablation also impacted more dorsal mossy cells that do not express calretinin (Houser et al., 2020; Botterill et al., 2021). In our initial data set (shown in Figure 1), the vast majority of mice demonstrated reliable viral targeting of mossy cells. After the initial data set was collected, we had a small number of instances in which our virus injections did not result in any cell transduction. Thus, to minimize the impact of technical factors on our ability to analyze the effects of mossy cell targeting in subsequent immunohistochemistry experiments, if slices did not demonstrate a >50% co-labeling with GFP of ventral calretinin-positive mossy cell bodies (for flex GFP or TeLC-GFP injected mice) or >50% loss of calretinin staining (taCasp3-injected Crlr-Cre mice relative to WT mice), mice were excluded from further analysis. These exclusion criteria only eliminated ∼15% of virus-targeted mice (reliably targeted mice for each group: GFP controls n = 42 of 47, MC ablated n = 43 of 49, and MC silenced n = 40 of 47); all Cre-negative (WT) mice injected with AAV5-flex-taCasp3 were included (n = 49 mice).

To assess dendritic morphology of retrovirus-labeled adult-born DGCs, confocal image stacks were taken with a 20X objective with 2X zoom (0.63µm step) or 40X oil objective (0.27µm step) for neurons at 14 dpi, and a 20X objective with no zoom for 21 dpi neurons. We collected a minimum of 3 cells per animal for dendritic tracing and no more than 5 cells from any individual mouse. These values were maintained as individual cellular values (i.e., n= cells instead of mice for these outcome measurements). Cells were excluded from dendritic analysis if proximal dendritic branches in the IML or MML were truncated due to sectioning. FIJI was used to measure the number of dendritic branches and total dendritic length for retrovirus labeled DGCs, based on manual tracing by a blinded observer. The Sholl analysis plugin in FIJI was used to quantify dendritic branching of adult-born DGCs.

Dendritic spines on 21 day-old retrovirus labeled DGCs were imaged in the proximal and distal regions of the molecular layer (IML and OML), defined as the innermost and outermost 40µm, using a 40X objective (8X zoom, 0.2µm step). For each cell, 2-4 dendritic segments were imaged and quantified in each region to obtain an average dendritic spine density for that cell (i.e., n= cells), using FIJI’s ROI manager. Granule cell axon (mossy fiber) sprouting was assessed in ZnT3-stained tissue, using z-stacks (10-15µm depth) with 20X and 40X objectives and 2 sections (one dorsal and one ventral) per mouse. Scoring of mossy fiber sprouting used a semi-quantitative blinded analysis with a score of 0 being no ZnT3 puncta in the granule cell layer, 1 for sparse ZnT3-positive puncta in the granule cell layer, 2 when ZnT3 puncta was regularly observed in the granule cell layer with occasional clusters in the proximal molecular layer, and a score of 3 being given to tissue that had ZnT3 puncta throughout the granule cell layer and a dense band of ZnT3 puncta in the proximal molecular layer, similar to previous reports (Shibley and Smith, 2002; Hunt et al., 2009). For VGlut2 and calretinin staining of fiber tracts in the molecular layer (perforant path and mossy cell axons, respectively), single stack images were taken with a 40X objective from the upper and lower blade of two tissue sections per mouse (one dorsal and one ventral) to assay the width of each of the relevant layers. All image analysis was performed by an experimenter blinded to experimental condition through the coding of file names prior to analysis.

### Electrophysiology

After terminal avertin anesthesia, mice were transcardiacally perfused with a choline chloride-based cutting solution containing the following (in mM): 110 choline chloride, 10 D-glucose, 1.3 Na-ascorbate, 25 NaHCO3, 7 MgCl2, 2.4 KCl, 1.25 NaH2PO4 dihydrate, and 0.5 CaCl2 with osmolarity balanced to ∼320mM and bubbled with 95% O2/5%CO2. Brains were blocked and glued to a sectioning stage for coronal sectioning at 300µm thickness using a Leica VT1200 vibratome. Slices were initially transferred to a storage chamber containing warmed (32-34oC) ACSF containing (in mM): 125 NaCl, 25 D-glucose, 25 NaHCO3, 1.25 NaH2PO4 dihydrate, 3 KCl, 1 MgCl2, and 2 CaCl2 with osmolarity balanced to 300-305mM and bubbled with 95% O2/5%CO2. After 30-45 minutes, slices were transferred to room temperature for (at least) an additional 30 minutes, before being placed in a submerged recording chamber (Siskiyou PC-H) on an upright, fixed-stage microscope equipped with infrared, differential interference contrast optics (Olympus BX50WI). Slices were continuously superfused with ACSF at 2 mL/min. Recordings were performed at room temperature from visually identified DGCs in the outer third of the granule cell layer (to target mature granule cells) or retrovirus-labeled adult-born DGCs (identified based on fluorophore expression, using fluorescence microscopy). To confirm proper viral targeting, acute slices were examined for GFP fluorescence in the IML prior to experimentation (for GFP or TeLC-GFP-infected tissue) or stained post-hoc to confirm the loss of calretinin-positive axons in the IML. In 1 of 5 AAV5-flex-GFP infected mice and 3 of 9 cases for AAV5 flex TeLC-infected mice, GFP expression was not observed and all slices were discarded prior to experimentation. In 2 of 8 cases for AAV taCasp3-infected mice, electrophysiology data were discarded based on ≥20µm calretinin-positive axons in proximal molecular layer in post-fixed slices.

Putative mature DGCs (unlabeled neurons recorded from the outer third of the granule cell layer) had lower input resistances than 21 day old retrovirus-labeled adult born DGCs (mature DGC= 253.7±17.7 MΩ n=23 cells, adult born DGC= 874.1±103.4 MΩ, n=16 cells; Student’s unpaired t-test, F(15,22)=18.23, p<0.0001) as is typical for adult-born DGCs (Overstreet Wadiche et al., 2005; Ge et al., 2007). There was no significant difference in neuronal input resistances between granule cell populations from different experimental conditions (mature cells: virus control= 230.8 ±26.3 MΩ n=9 cells, MC ablated= 255.4±33.1 MΩ n=7 cells, MC silenced= 284.9 ±35.5MΩ n=7 cells; One Way ANOVA, F(2,20)=0.009, p=0.79; adult born cells: virus control= 845.2 ±216.8 MΩ n=5 cells, MC ablated= 899.6.4 ±161.3 MΩ n=6 cells, MC silenced= 708.0 ±65.8MΩ n=5 cells; One Way ANOVA, F(2,13)=2.053, p=0.37).

Recording pipettes for non-fluorescent granule cells were pulled from borosilicate glass (TW150F; World Precision Instruments; Claremont, CA) with a P-87 puller (Sutter Instruments, Novato, CA), while recording pipettes for fluorescent (adult-born) DGCs were pulled from borosilicate leaded glass (PG10165; World Precision Instruments; Claremont, CA) with a PC-10 puller (Narishige International; Amityville, NY) to facilitate higher seal resistances. The intracellular solution for voltage clamp recordings contained (in mM): 113 CsGluconate, 8 NaCl, 10 EGTA, 10 HEPES, 1 MgCl2, 1 CaCl2, 3 CsOH, 0.3 MgATP and 2 NaGTP; pH 7.3. Pipette open tip series resistances were 5-7 MΩ. For recordings from adult-born granule cells, Alexa594 or Alexa488 were included in the pipette solution of a subset of neurons, to visually confirm patching/filling of the correct cell. Recordings were obtained using an Axopatch 1D amplifier (Molecular Devices, Sunnyvale, CA), low-pass filtered at 5 kHz, digitized at 10 kHz with a NIDAQ (National Instruments) analog-to-digital board, and acquired using Igor Pro (Wavemetrics) software with NIDAQmx (National Instruments) plugins.

Cells were whole-cell voltage-clamped at -70mV for 5-10 min to allow equilibration of pipette and intracellular solutions prior to collection of evoked responses. Cell series and input resistances were measured using a -10mV voltage step (50 msec) prior to each sweep. The input resistance was calculated using V=IR, using the plateau (steady-state) current required to hold the -10mV step after resolution of capacitive currents. To evoke currents from proximal molecular layer inputs, a bipolar electrical stimulating electrode (FHC; Bowdoin, ME) was placed in the inner molecular layer ∼20µm above the edge of the granule cell layer and ∼100-150µm from the recorded neuron, and stimulating pulses were delivered by an ISOflex stimulus isolation unit (A.M.P.I.). Stimulation intensity was adjusted for each cell to deliver an evoked synaptic response of ∼200pA. The GABAA-receptor antagonist SR95531 (10µM) was applied to isolate excitatory synaptic inputs during electrical stimulation when indicated. To assess the ratio of excitatory:inhibitory synaptic responses, the amplitude of the pharmacologically-isolated evoked EPSC (after SR95531) was divided by the amplitude of the SR95531-sensitive (GABAaR-mediated) inhibitory evoked current, obtained by subtraction. After a steady-state excitatory response was maintained for a minimum of ∼4 minutes, in some experiments we applied the cannabinoid receptor-1 (CB1R) agonist Win 55,212 (5µM).

### Acute seizure modeling

We assessed acute seizure susceptibility in adult mice 6 weeks after AAV injection as previously described (Danis et al., 2024). Pentylenetetrazol (PTZ; Sigma) was diluted in sterile saline at 4 mg/mL and administered to a subset of mice by intraperitoneal injection at a dosage of 40 mg/kg. Mice were scored for seizures for 1 hour after PTZ administration, once per minute using a modified Racine scale for mice (Shibley and Smith, 2002), from zero for normal behavior to six for high-amplitude jumping/popcorn behavior. The score for each minute reflected the highest-grade seizure observed in the prior minute. To obtain a cumulative seizure score for each mouse, we added the maximum seizure score observed for each minute of the first 20 minutes following saline or PTZ administration. Seizure onset was defined as the post- injection timepoint when the mouse first exhibited a score of 3 (unilateral forelimb myoclonus) or greater. One hour after PTZ injection, mice were humanely euthanized for brain harvest and subsequent staining for assessment of cellular activity.

### Data Analysis

Graphpad Prism software was used for statistical analysis. All data was assessed for normal distribution using Shapiro-Wilk normality testing. For normally distributed data, a Student’s unpaired t-test was used to assess differences between the appropriate virus control and mossy cell-manipulated mice. For immune response data a two-way ANOVA with Sidak’s post-doc tests were used. Molecular layer axon staining measurements (layer widths) and functional responses to electrical stimulation of the IML GFP were not significantly different from the two control groups (Cre-negative Crlr-Cre mice injected with AAV5-flex-taCasp3 and Cre-positive mice injected with AAV5-flex-GFP) and were combined into a single control group on graphs for ease of presentation. Data from the combined control group was compared to silenced or ablated mice using one-way ANOVAs with Tukey’s multiple comparisons. When data were not normally distributed, non-parametric testing was used. Significance was set at p<0.05. Data are presented as mean ±SEM, and these values as well as their respective statistical analyses are listed in Table 1 if not mentioned in the results.

**Table 1:**
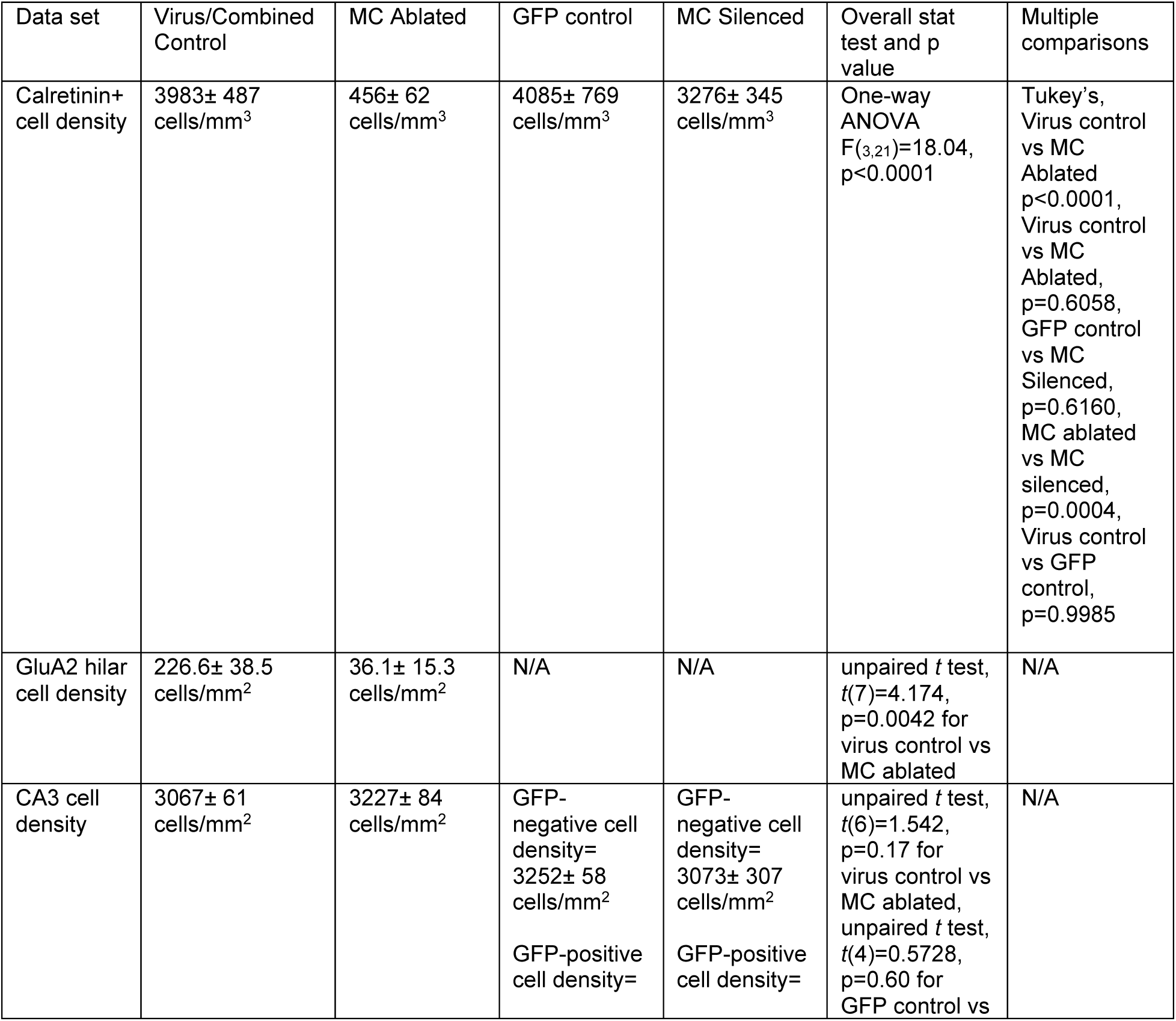
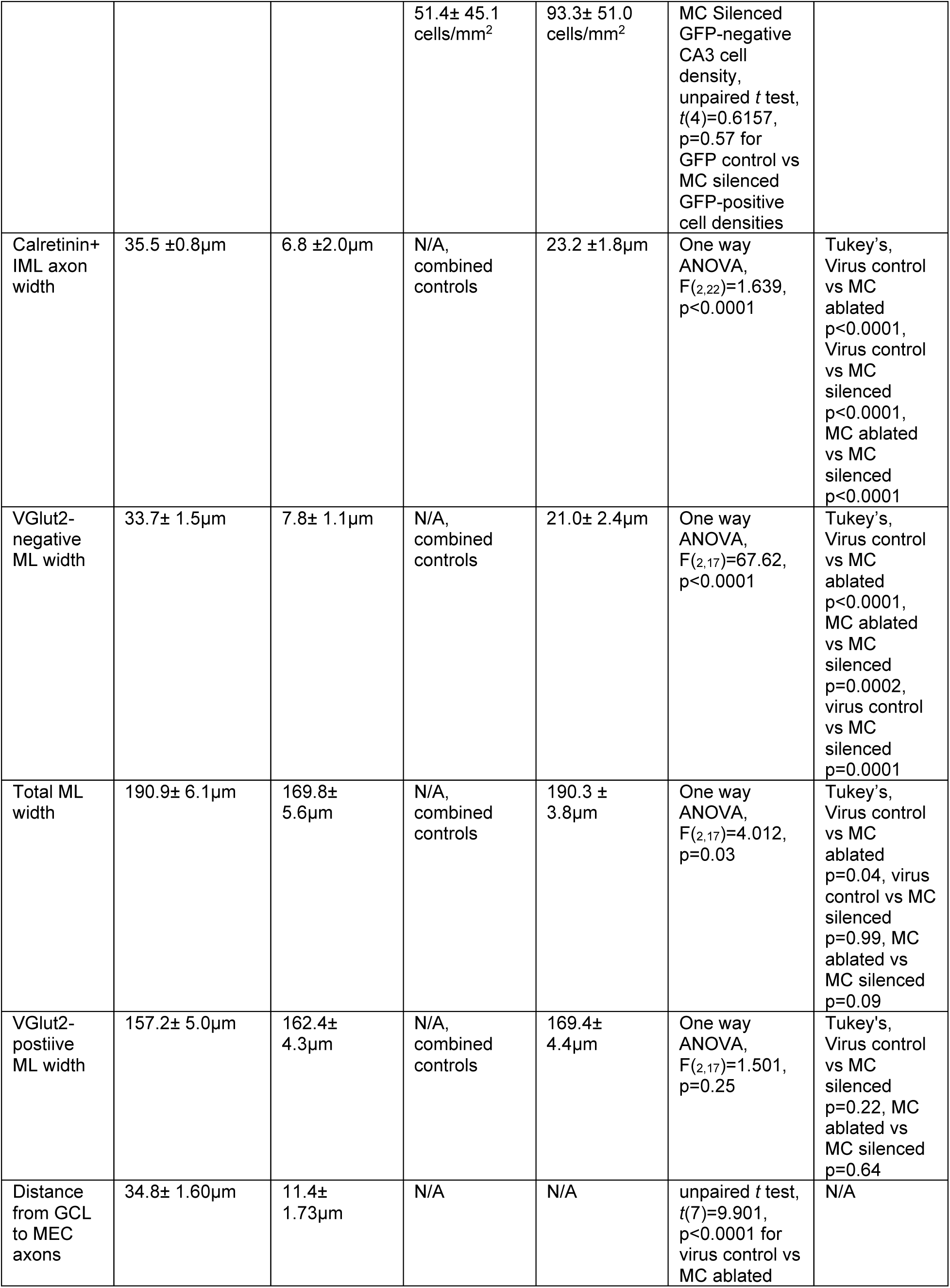
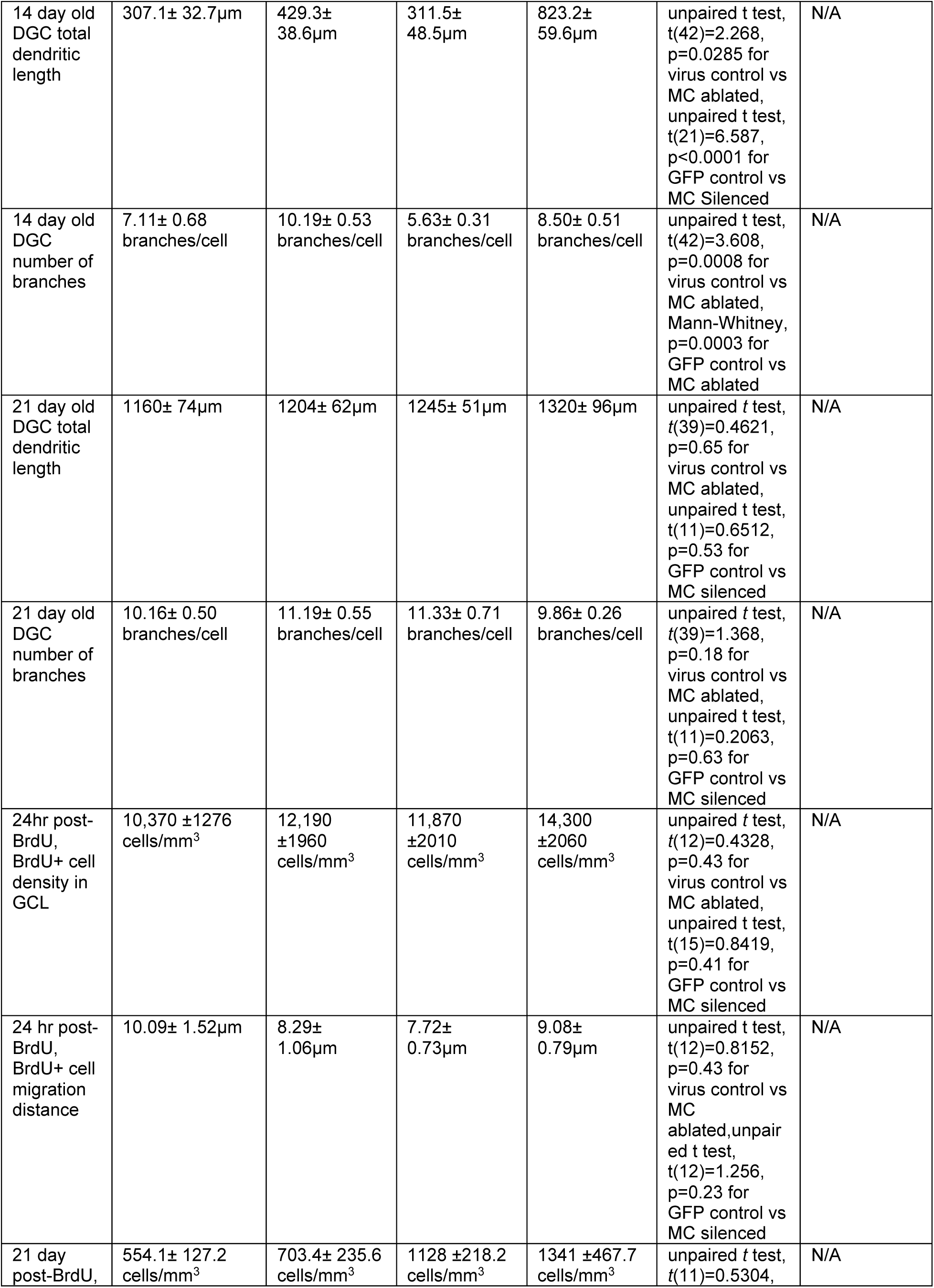
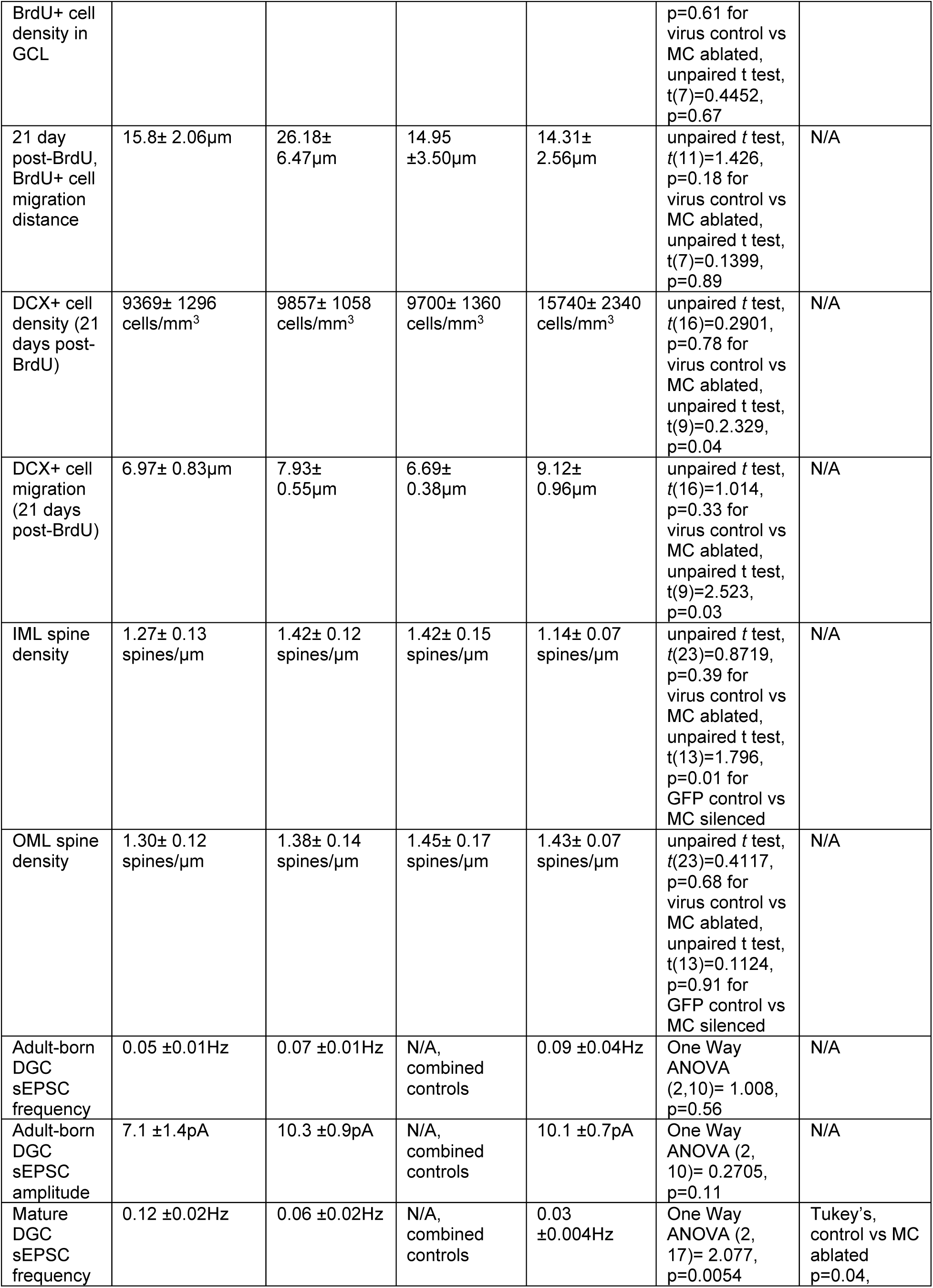
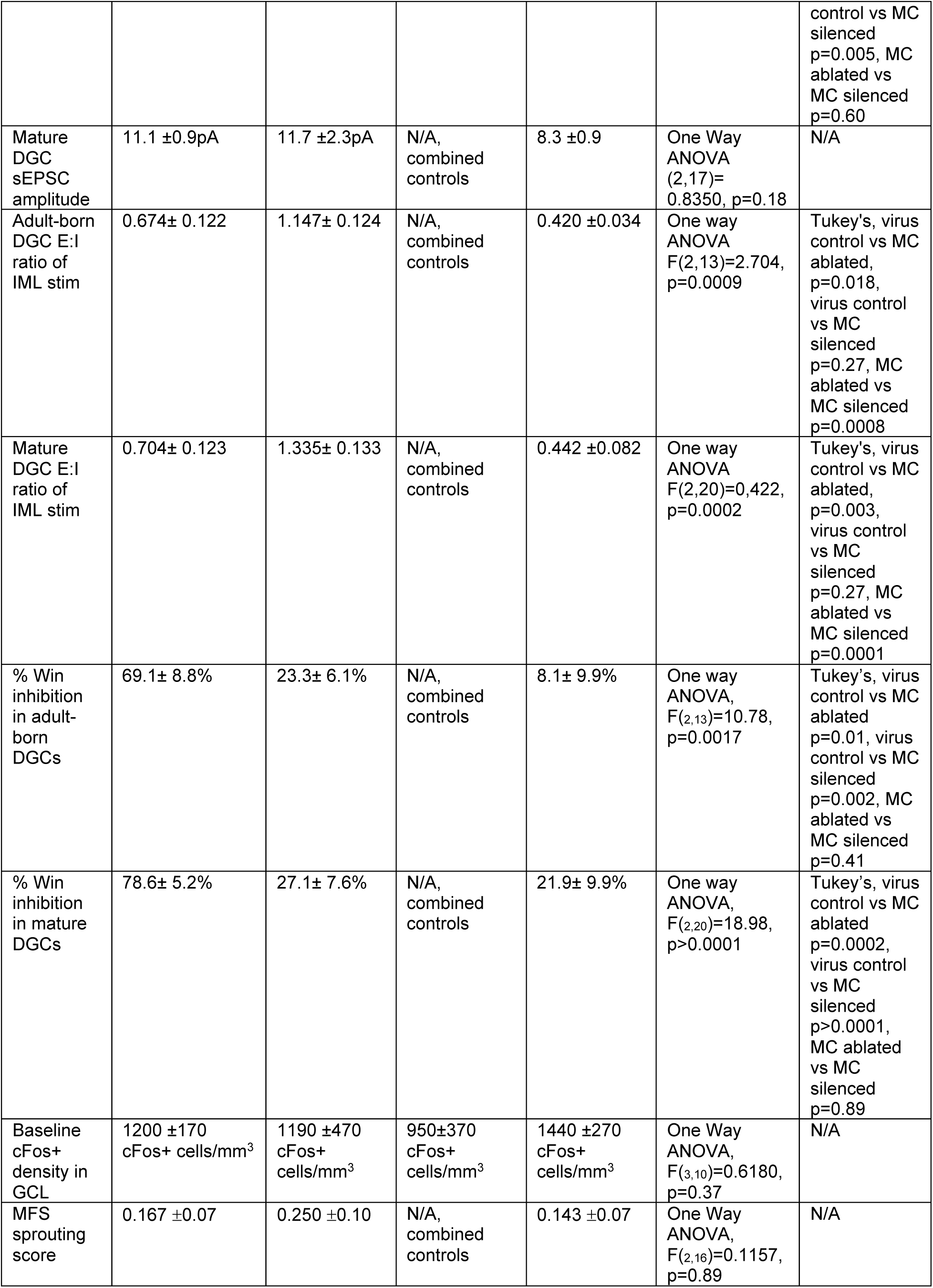
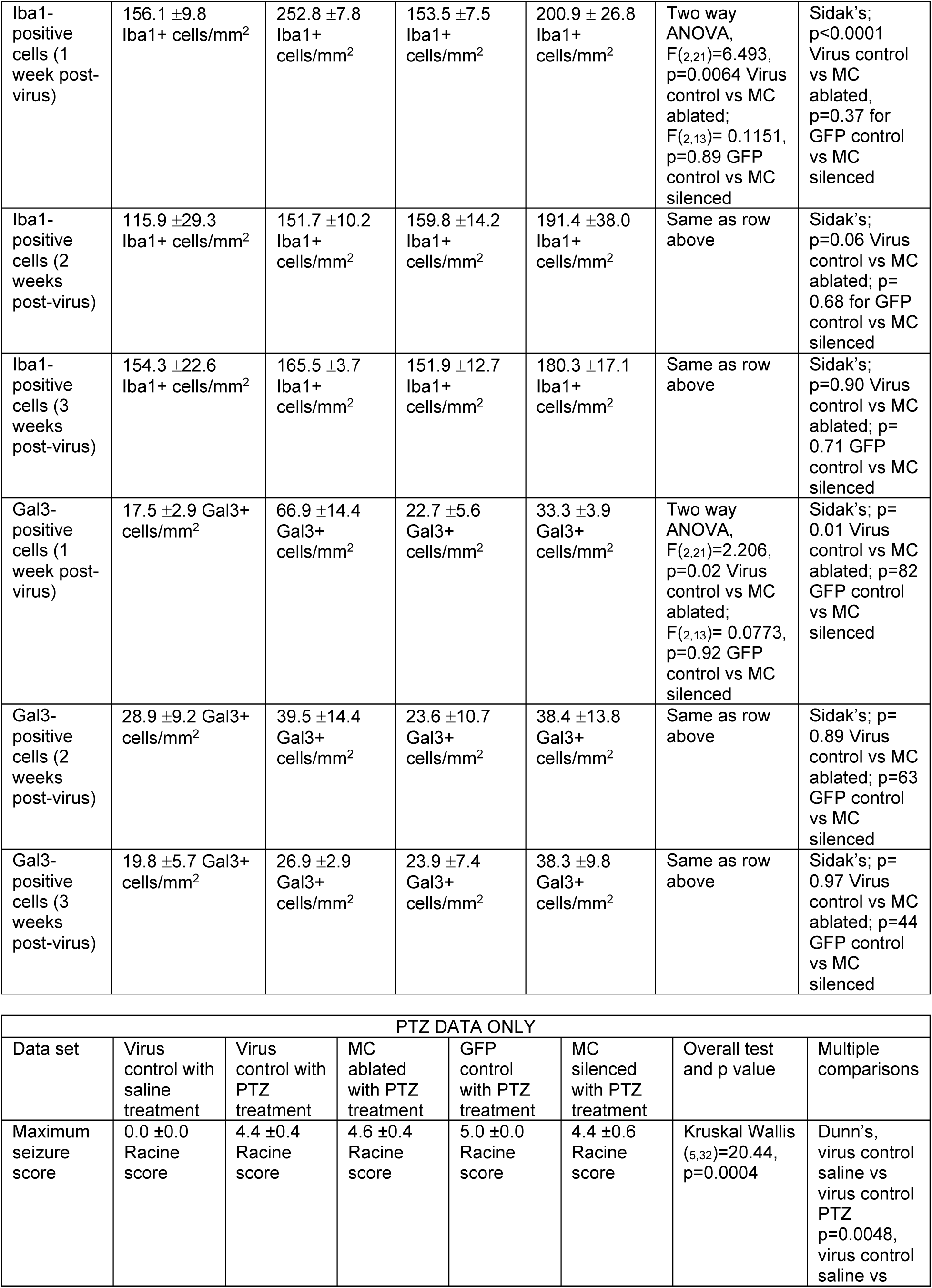
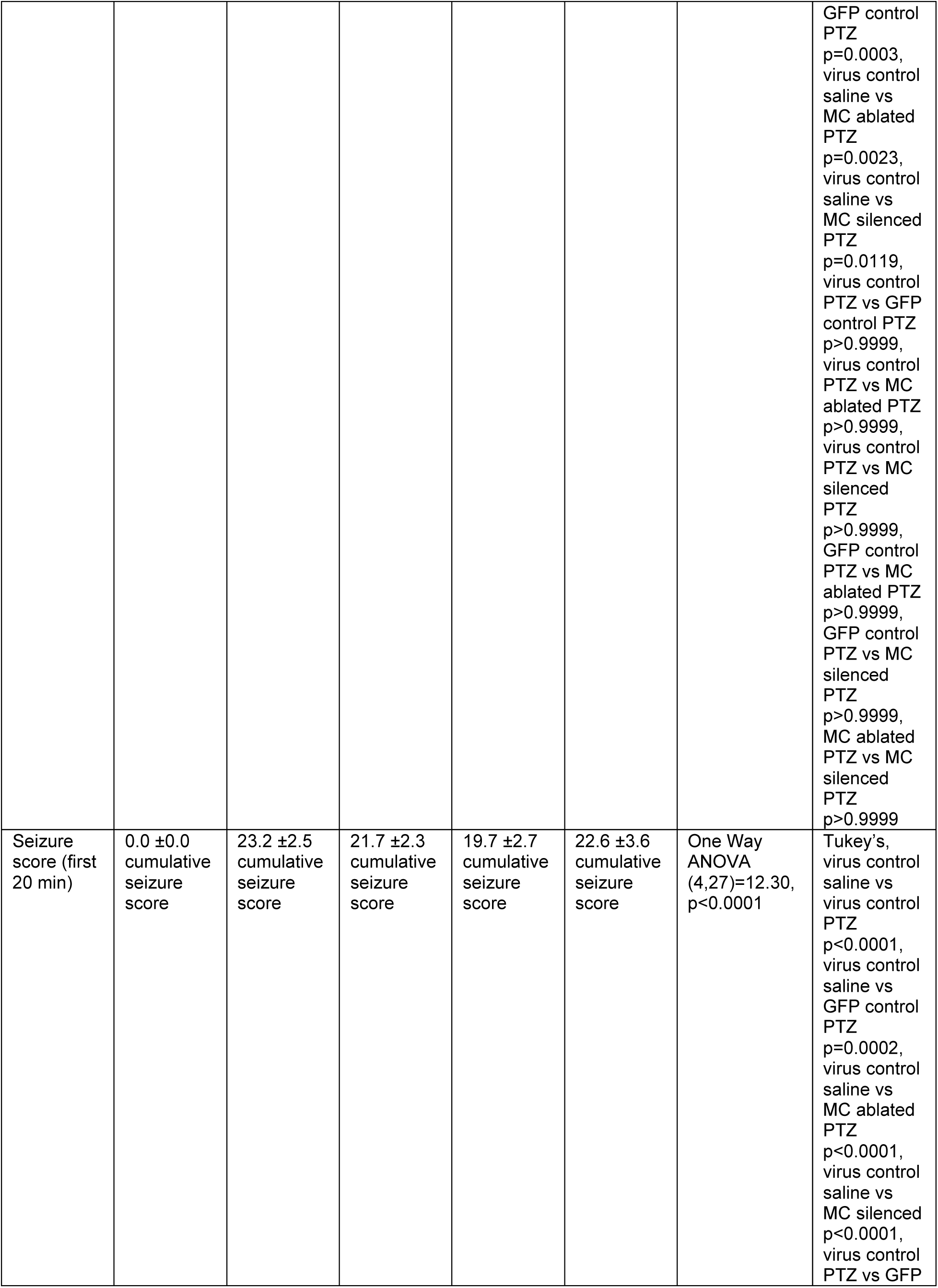
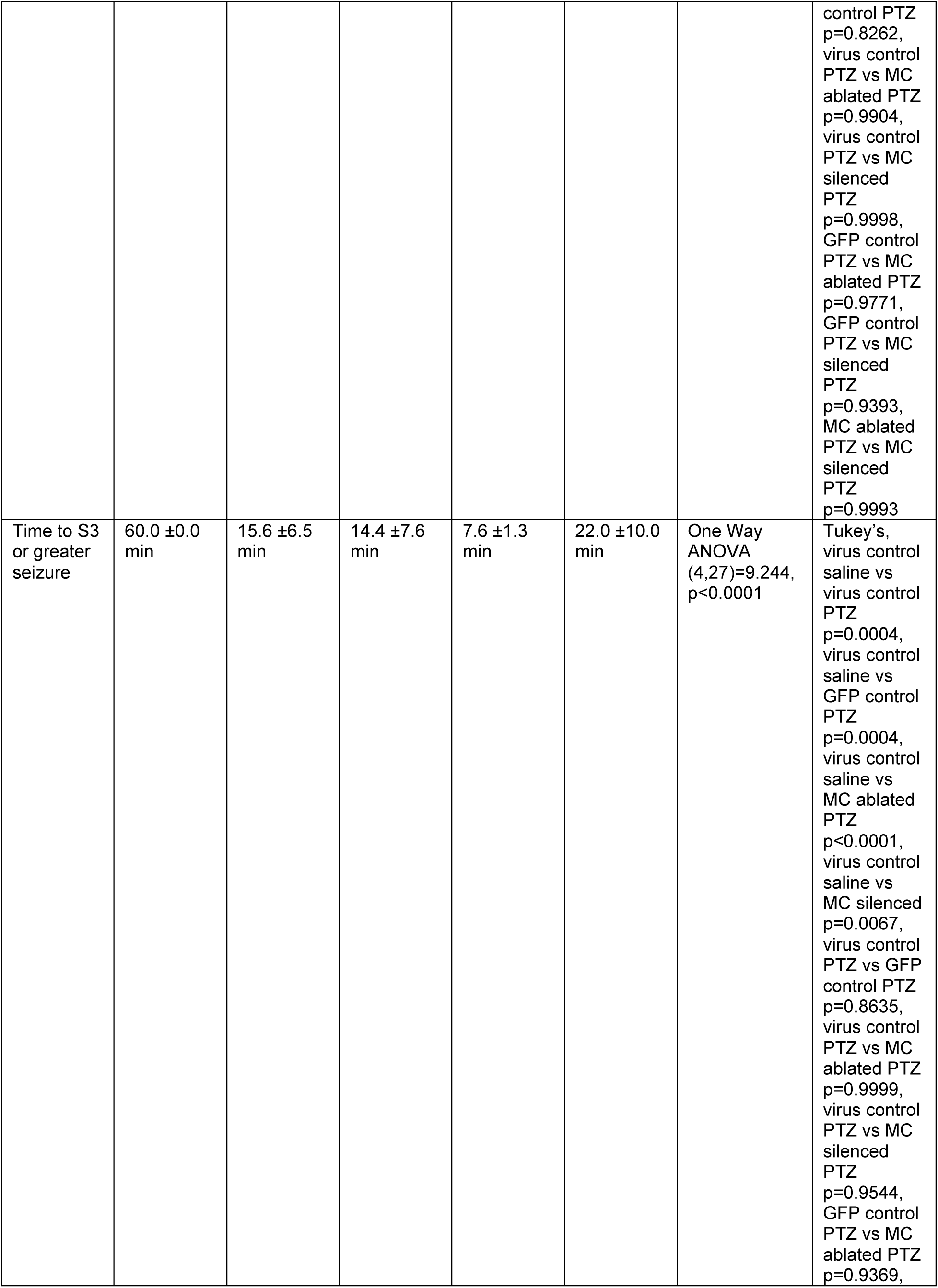
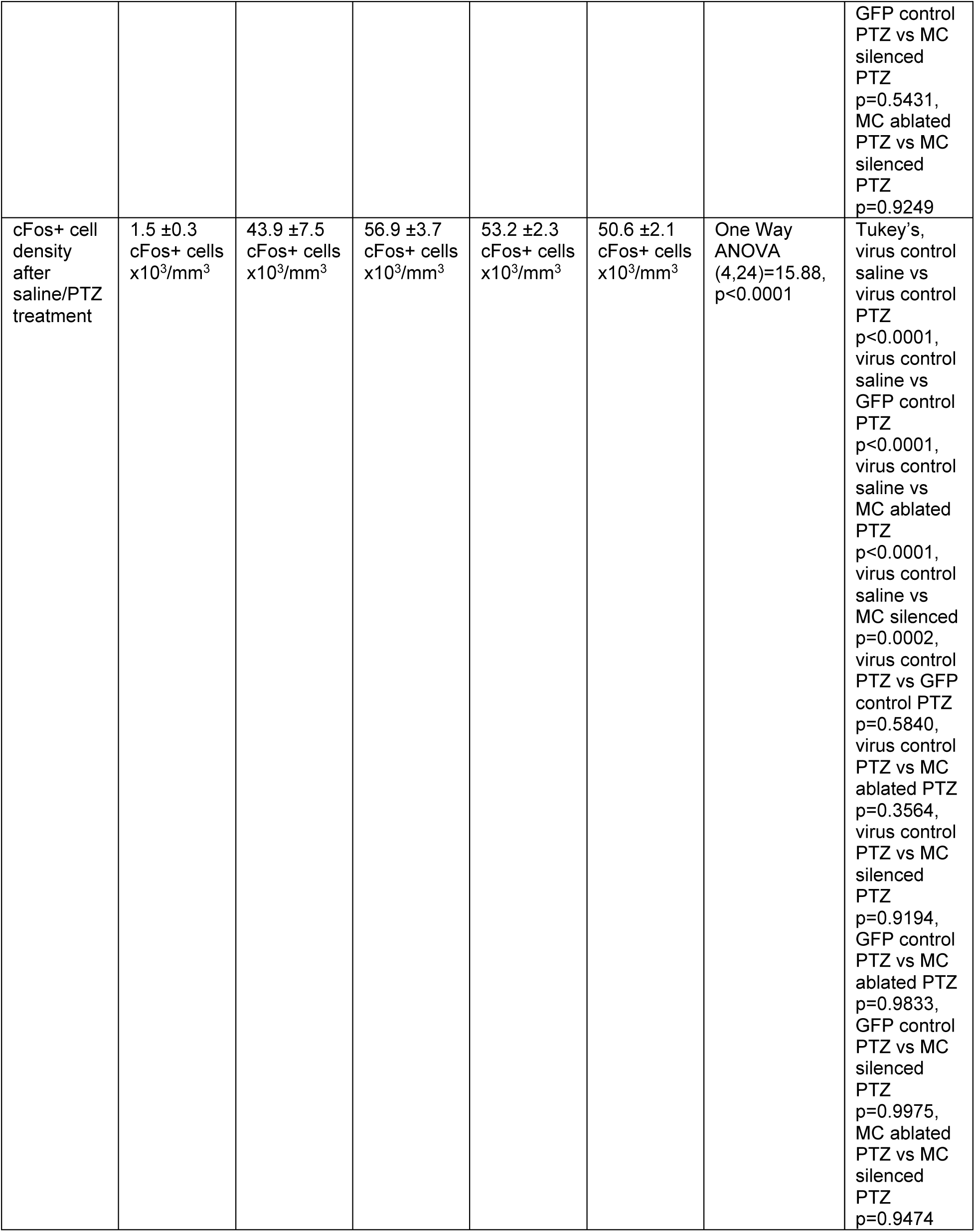
Summary data and statistical comparisons used in the Results.

## Acknowledgements

We thank Dr. Hiro Nakazawa for providing mutant mice via Jackson Laboratories, Drs. Peer Wulff and Nirao Shah for viral vector constructs, Daniel Kim for technical support, the OHSU Advanced Light Microscopy Core (RRID:SCR_009961) for expert technical assistance, and members of the Schnell and Westbrook labs for feedback and support. This work was funded by NIH Grants R01NS126247 (ES), R21NS102948 (Ines Koerner / ES), R01NS117371 (GLW), NIH Grant P30-NS061800 (OHSU Advanced Light Microscopy Core), F32NS106732 (CRB), VA I01-BX004938 (ES), VA I01-BX006921 (ES), and VA IK2-BX005761 (CRB). The contents of this manuscript do not represent the views of the US Department of Veterans Affairs or the US government.

---

**Supplemental Figure 1:**
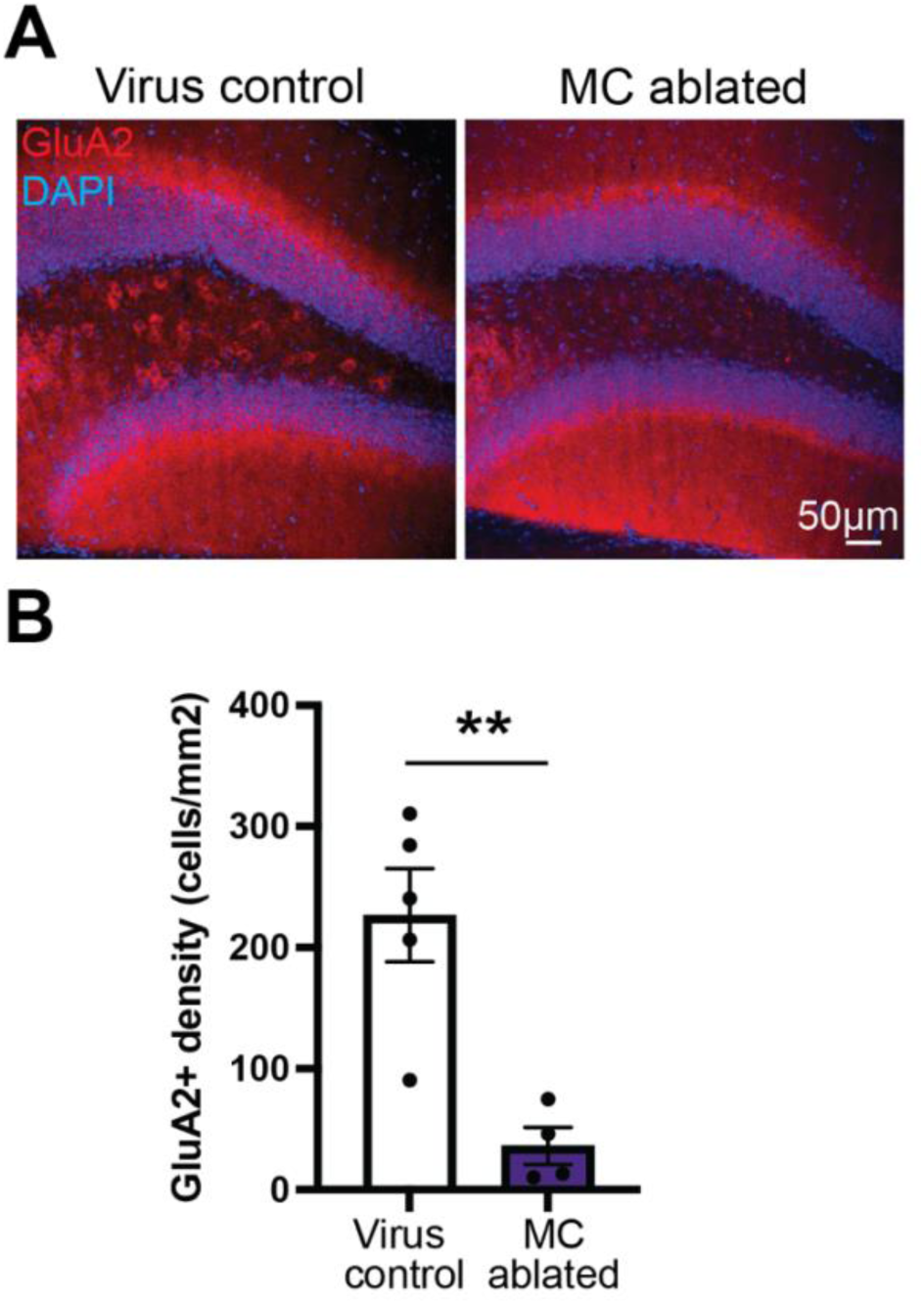
Mossy cell ablation by AAV5-flex-taCasp3 in dorsal hippocampus. A) Representative sections from dorsal hippocampus from virus control and mossy cell (MC) ablated mice, 6 weeks after AAV5-flex-taCasp3 virus injection. Tissue was stained for glutamate receptor AMPA type subunit 2 (GluA2, red) and co-stained for DAPI (blue) to visualize the dentate granule cell layer and CA3 pyramidal cell layer, which also express GluA2. B) GluA2+ hilar cell density is reduced in MC ablated (N=4 mice) mice compared to virus control (N=5 mice) mice (** p>0.01).

**Supplemental Figure 2:**
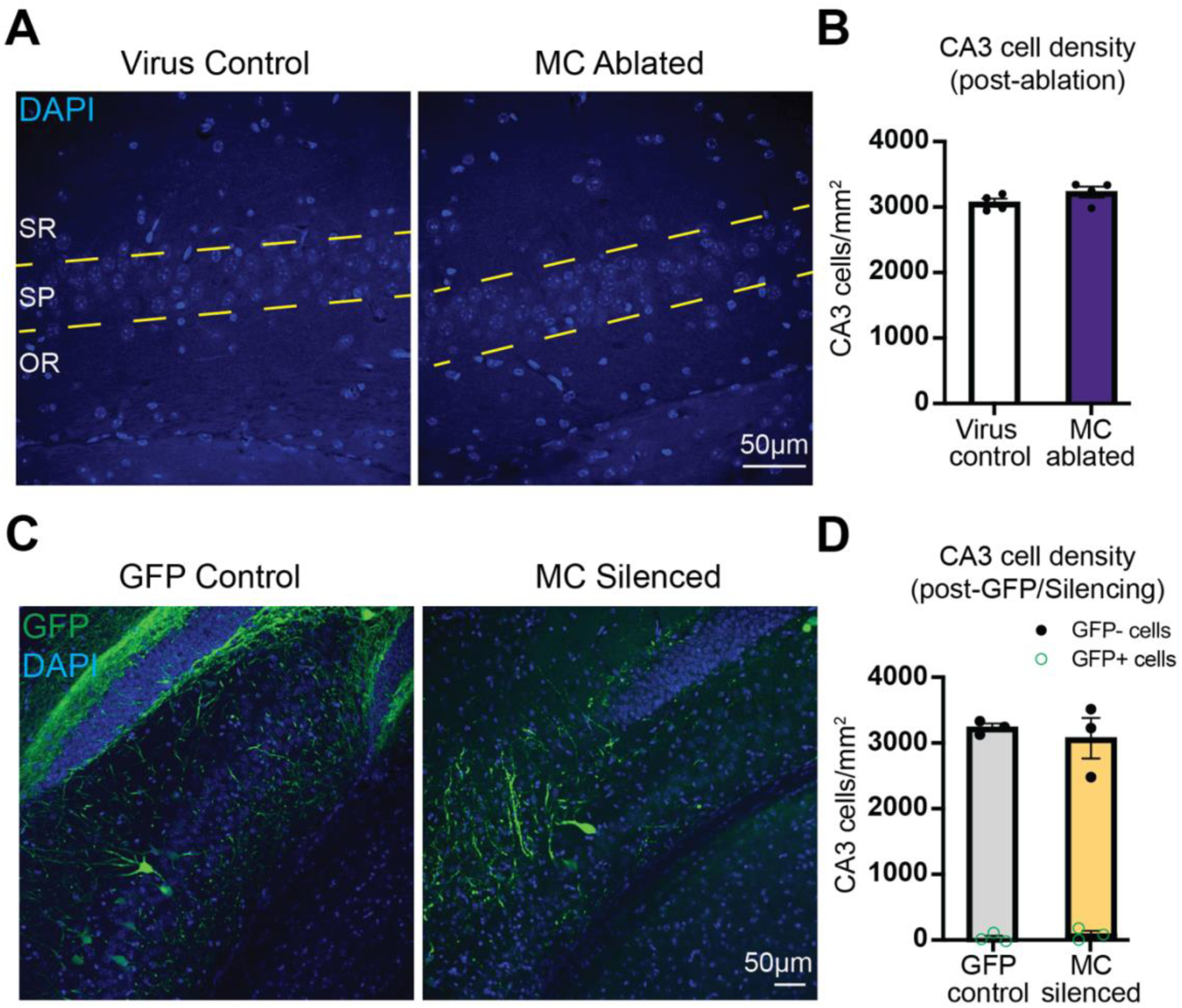
Minimal off-target viral expression in CA3 neurons. A) Representative images of DAPI-stained pyramidal cell nuclei in proximal CA3 from virus control and mossy cell (MC) ablated mice. Yellow dashed lines represent borders of the pyramidal cell layer (SP), stratum oriens (OR) and stratum radiatum (SR). B) CA3 cell densities were similar between virus control (N= 4 mice) and MC ablated conditions (N= 4 mice), suggesting minimal ablation of CA3 pyramidal neurons following AAV5-flex-taCasp3 virus injection (p=0.17). C) Representative images of GFP-expressing neurons in CA3 from Crlr-Cre mice injected with AAV5-flex-GFP or AAV5-flex-TeLC-GFP viruses, co-stained with DAPI. D) Cell densities of both GFP-negative (GFP control, N= 3 mice; MC Silenced, N= 3 mice; p=0.60) and GFP-positive (p=0.57) were not different between experimental groups.

**Supplemental Figure 3:**
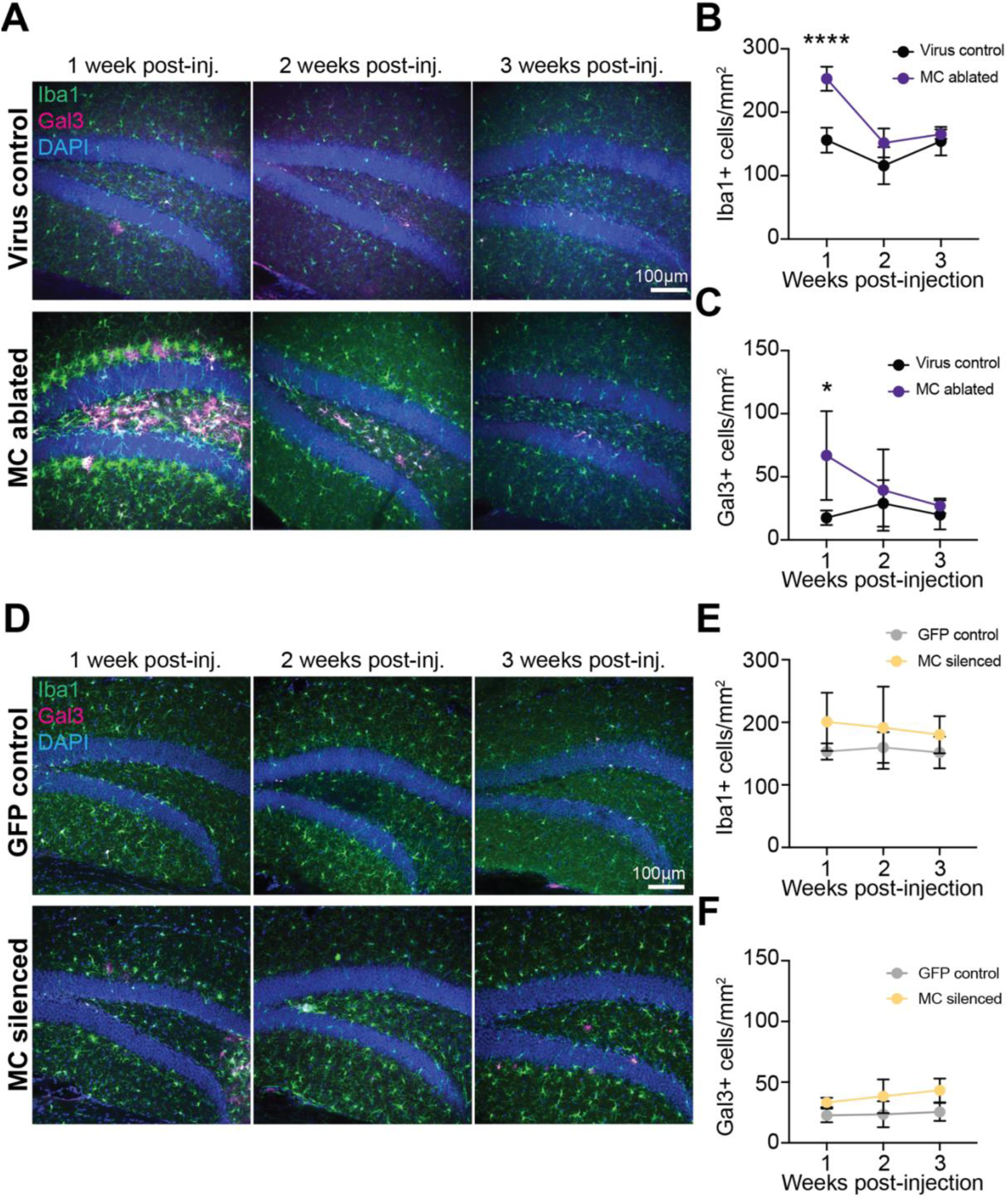
Timecourse of microglial activation after mossy cell ablation. A) Representative dentate sections 1, 2, or 3 weeks following AAV flex Casp3 injections into Crlr-Cre-negative (Virus control) or Crlr-Cre-positive (MC ablated) mice. Microglia are stained with anti-Iba1 (green), with activated microglia identified by anti-Gal3 (red) staining. B) Iba1-positive cell densities at different timepoints after virus injection; (Virus control n = 4 mice at 1 week post-virus, 4 mice at 2 weeks, and 4 mice at 3 weeks; MC ablated n = 6 mice at 1 week, 5 mice at 2 weeks, and 4 mice at 3 weeks (**** p <0.0001 at 1 week; n.s. at 2 and 3 weeks). C) Gal3-positive cell densities at different timepoints after virus injection (* p <0.05 at 1 week; n.s. at 2 and 3 weeks). D) Representative dentate sections 1, 2, and 3 weeks following AAV flex GFP (GFP control) or AAV flex TeLC-GFP (MC silenced) injections into Crlr-Cre-positive mice. Microglia are stained with anti-Iba1 (green, pseudocolored), with activated microglia identified using anti-Gal3 (red) staining. E) Iba1-positive cell densities at different timepoints after virus injection (GFP control n = 3 mice at 1 week post-virus, 3 mice at 2 weeks, and 4 mice at 3 weeks; MC silenced= 3 mice at 1 week, 3 mice at 2 weeks, and 3 mice at 3 weeks (n.s. at all timepoints). F) Gal3-positive cell densities at different timepoints after control virus injection or MC silencing (n.s. at all timepoints).

**Supplemental Figure 4:**
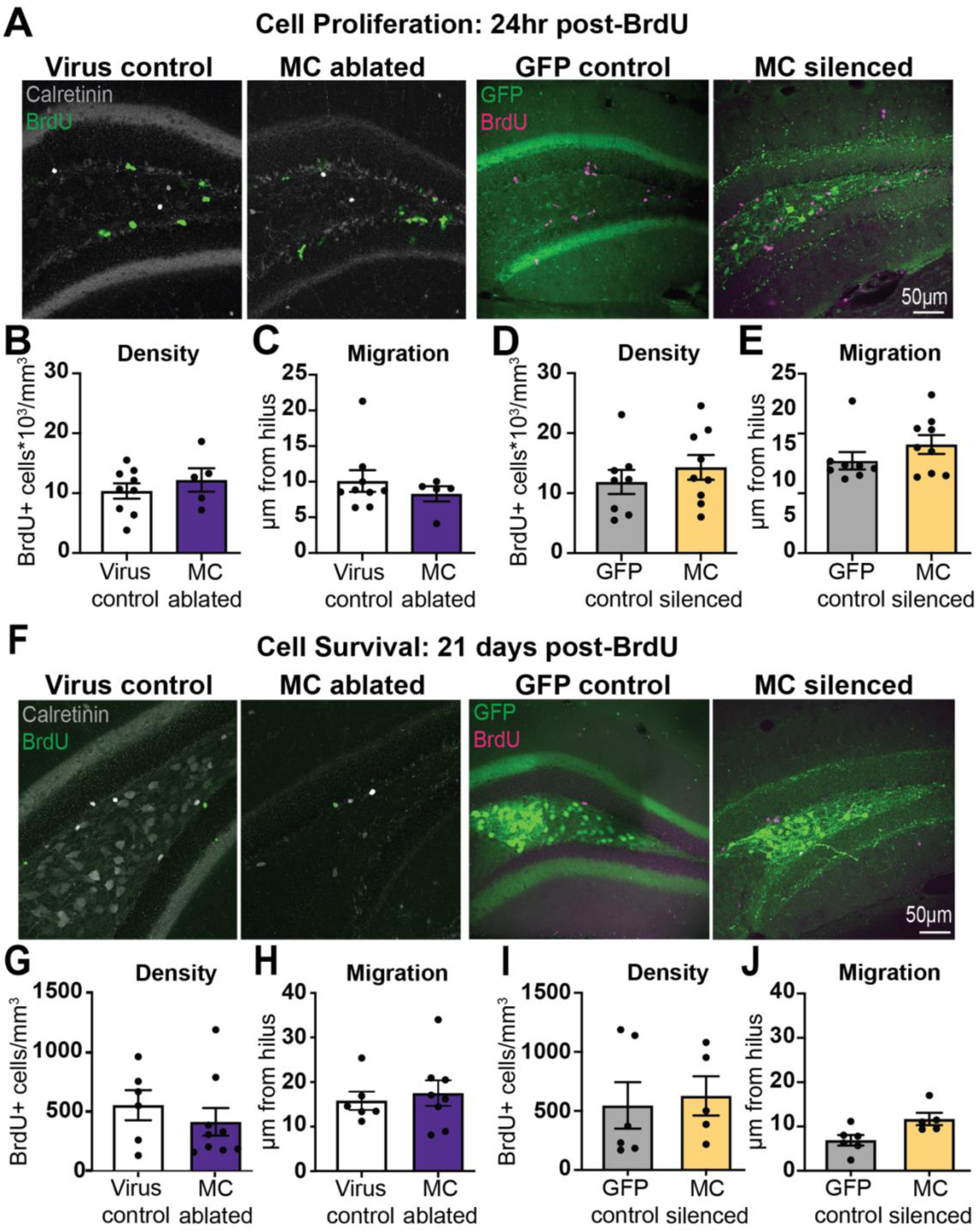
Cell proliferation and survival in the dentate granule cell layer are not altered following mossy cell manipulations. A) Representative mitotic labeling in virus controls, MC ablated, and MC silenced conditions 24hr after BrdU administration to assess cell proliferation. B&C) Cell proliferation and outward migration in virus control and MC ablated conditions (Virus control n = 9 mice, MC ablated n = 5 mice; n.s. both measures). D&E) Cell proliferation and outward migration in GFP control and MC silenced conditions (GFP control n = 8 mice, MC silenced n = 9 mice; n.s. both measures). F) Representative BrdU labeling in virus controls, MC ablated, and MC silenced conditions 21 days after BrdU administration to assess newborn cell survival. G&H) BrdU-positive cell density and outward migration distance in virus control and MC ablated conditions, 21 days after BrdU administration (Virus control n = 6 mice, MC ablated n = 9 mice; n.s. both measures). I&J) BrdU-positive cell density and outward migration distance in GFP control and MC silenced conditions, 21 days after BrdU administration (GFP control n = 6 mice, MC silenced n = 5 mice; n.s. both measures).

**Supplemental Figure 5:**
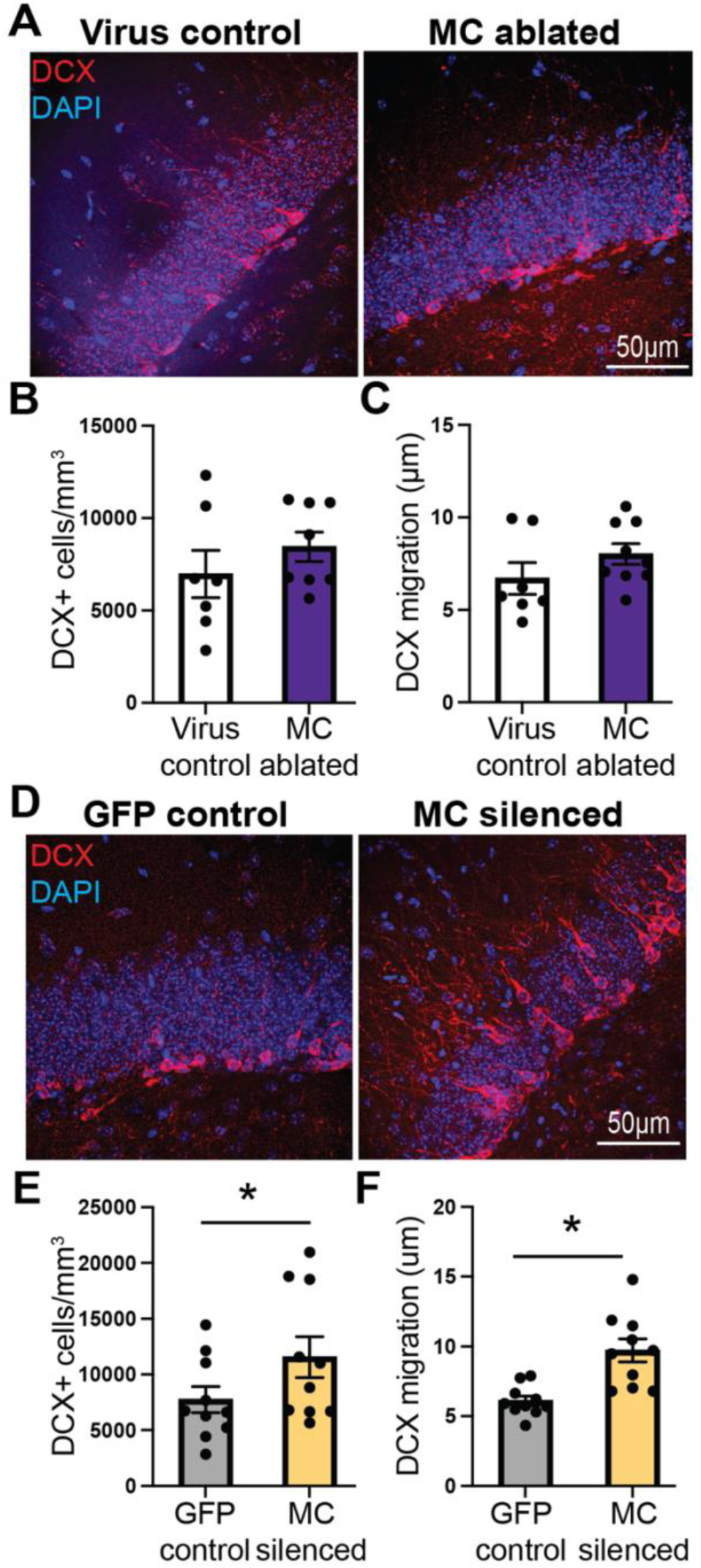
Differential effects of mossy cell ablation vs silencing on immature granule cells. A) Representative immature adult-born granule cells, as assessed by doublecortin (DCX) staining (red) in virus control and MC ablated mice. B&C) DCX-positive cell density and outward migration from the hilus, demonstrating no change following mossy cell ablation (Virus control n = 7 mice, MC ablated n = 8 mice; n.s. both measures). D) Representative immature adult-born granule cells (DCX+; red) in GFP control and MC silenced mice. E&F) DCX staining demonstrates a mildly increased density of immature granule cells and increased outward migration of immature cells following mossy cell silencing (GFP control n = 10 mice, MC silenced n = 10 mice; * = p <0.05).

**Supplemental Figure 6:**
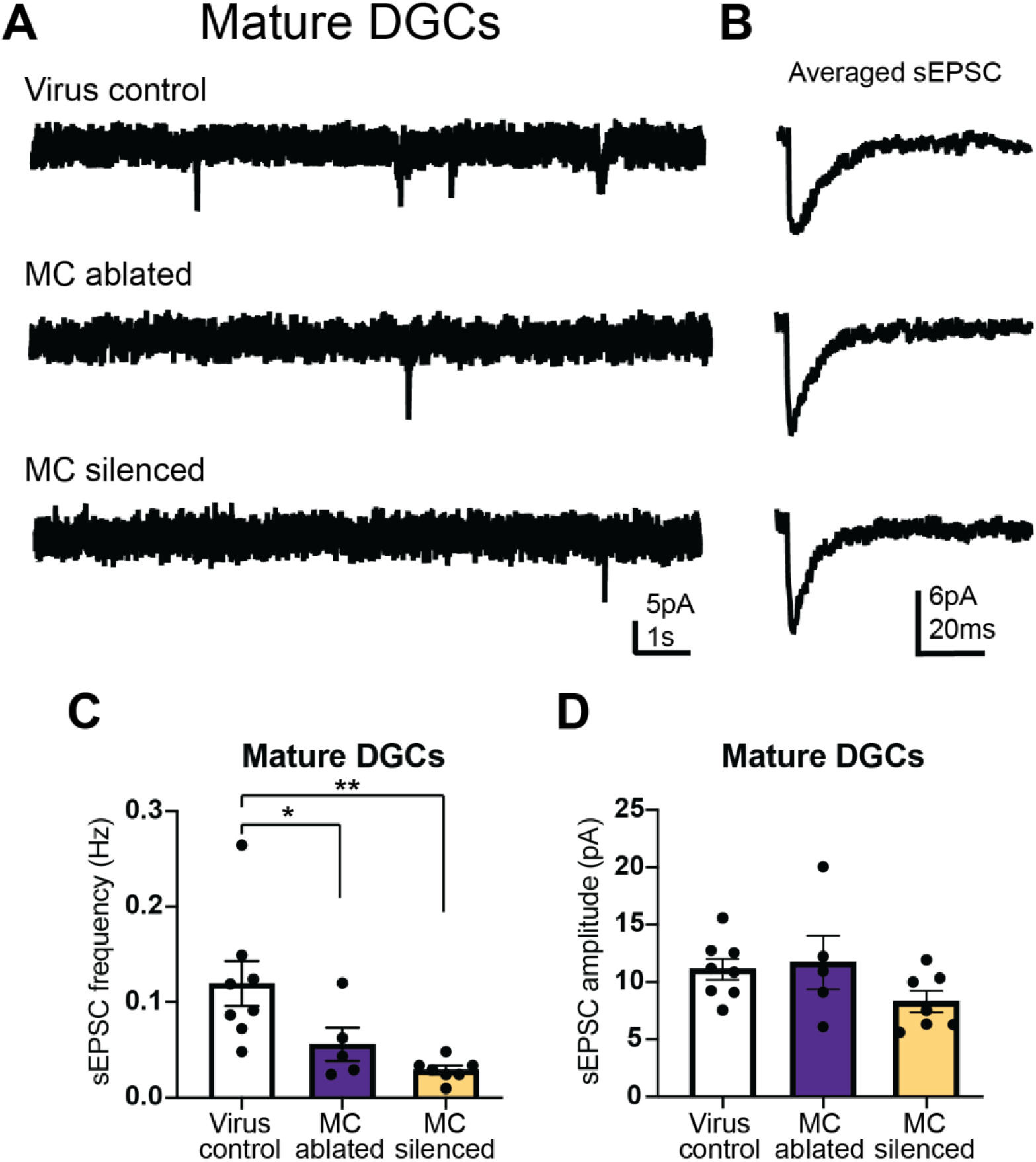
Reduced spontaneous excitatory synaptic currents in mature DGCs after mossy cell ablation or silencing. A) Representative spontaneous excitatory post-synaptic current (sEPSC) recordings from mature DGCs from virus control, MC ablated, and MC silenced mice. B) Average sEPSC waveforms from the respective recordings in A. C&D) sEPSC frequencies and amplitudes recorded from mature DGCs from virus control, MC ablated, and MC silenced mice. (sEPSC frequency * = p < 0.05, ** p <0.01; sEPSC amplitude n.s. all comparisons).

**Supplemental Figure 7:**
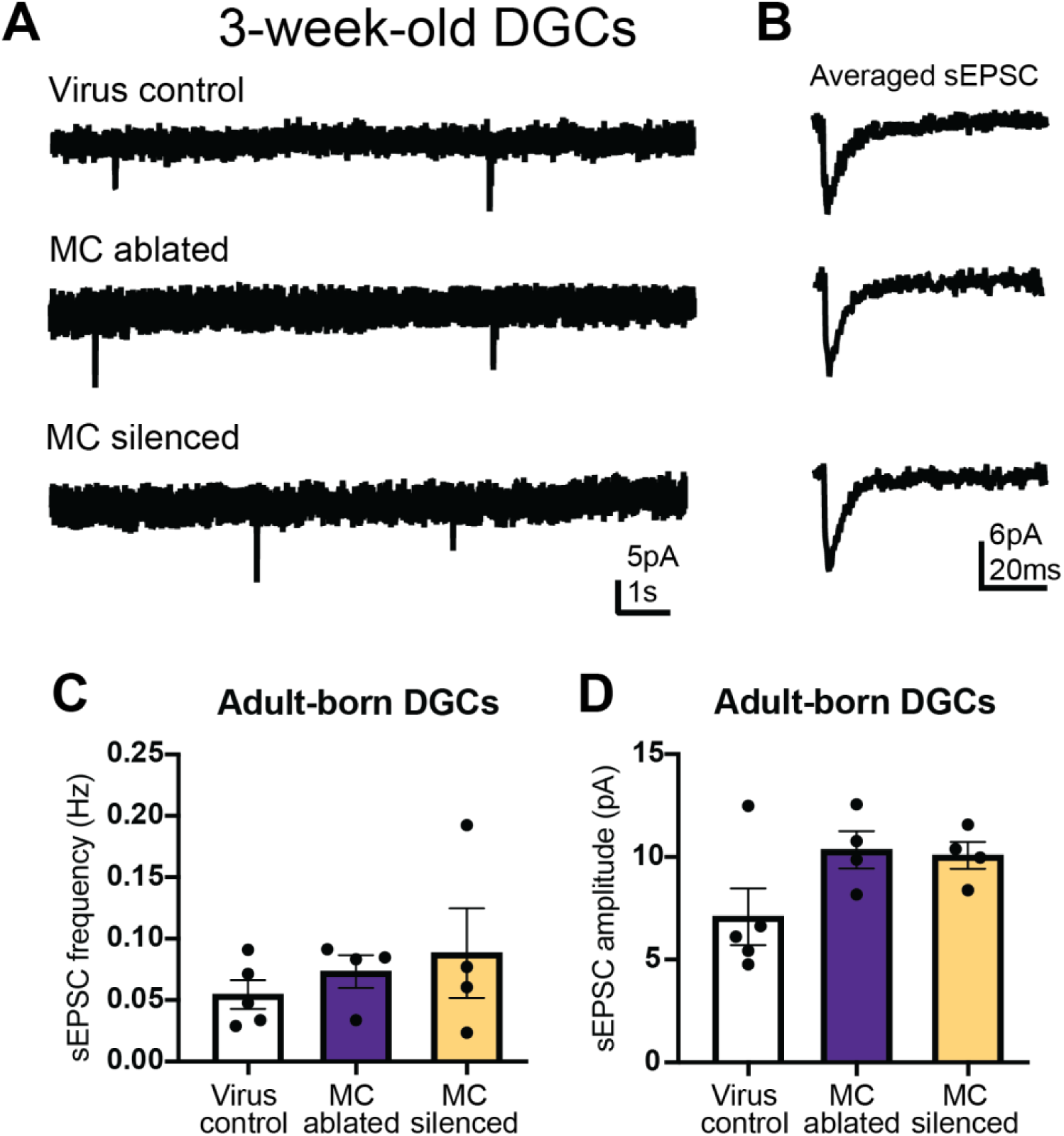
sEPSCs are unchanged in immature DGCs generated after mossy cell ablation or silencing. A) Representative sEPSC recordings from 21 day-old adult-born DGCs from virus control, MC ablated, and MC silenced mice. B) Average sEPSC waveforms from the respective recordings in A. C&D) sEPSC frequencies and amplitudes in 21 day old DGCs from virus control, MC ablated, and MC silenced mice. Neither sEPSC frequency or amplitude was different in adult-born cells in each group. (n.s. all comparisons).

**Supplemental Figure 8:**
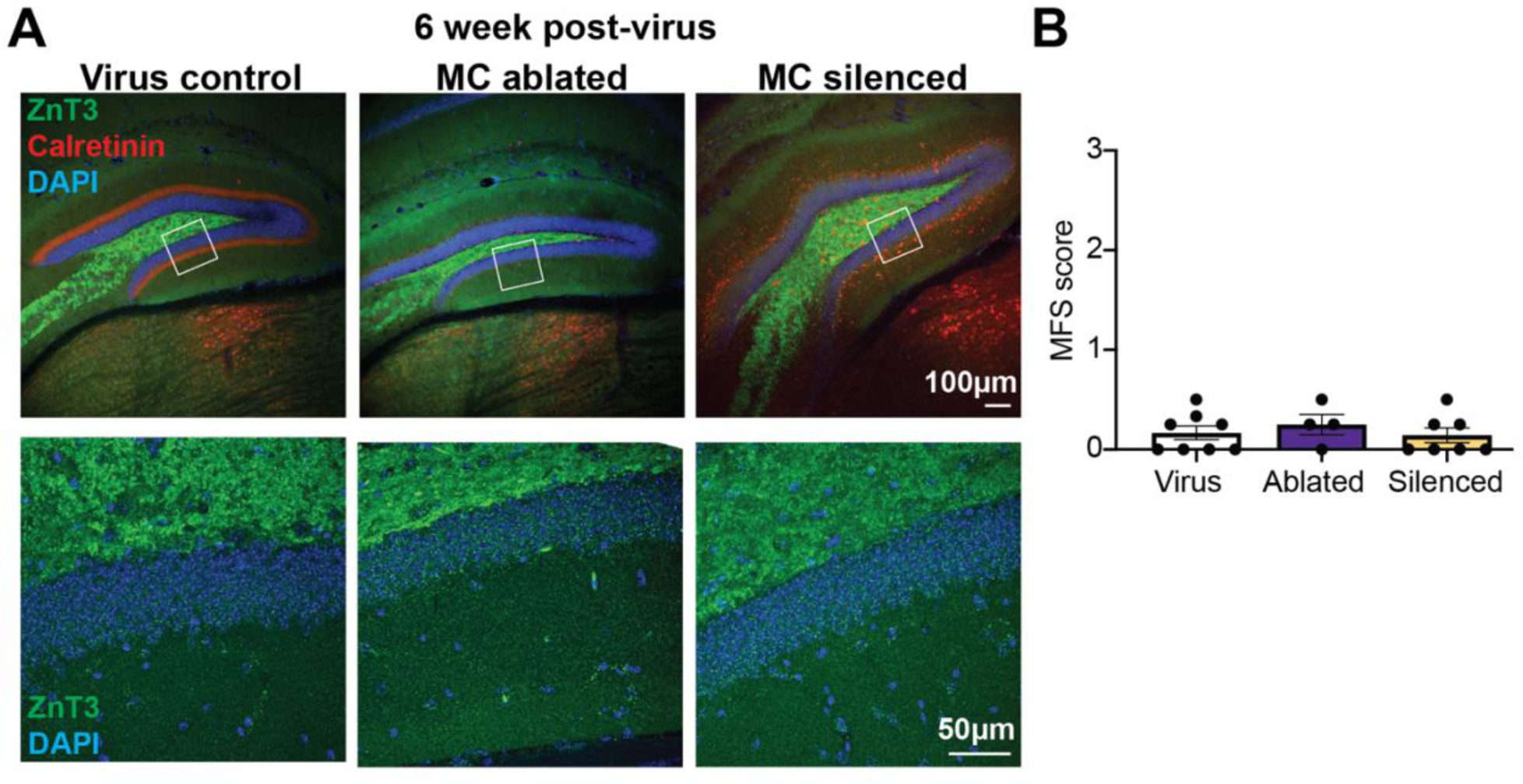
Neither mossy cell ablation or silencing drive recurrent granule cell axon (mossy fiber) sprouting. A) Representative 5X (top) and 40X images (bottom) of mossy cell axon (calretinin, red) and granule cell mossy fiber bouton (ZnT3, green) staining in virus control, MC ablated, and MC silenced conditions 6 weeks after virus injection. B) Mossy fiber sprouting (MFS) is not detected in either MC ablated or MC silenced groups (Virus control n = 8 mice, MC ablated n = 4 mice, MC silenced n = 8 mice; n.s. all groups).

**Supplemental Figure 9:**
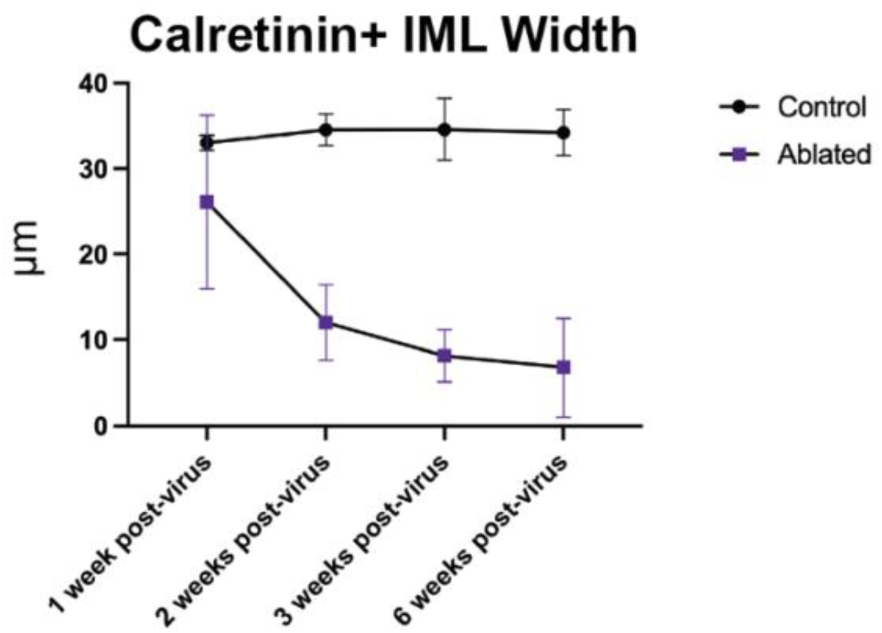
Calretinin-positive IML width decreases during the first 2 weeks after mossy cell ablation and remains stable at later time points.

